# MAGIC: A diffusion-based imputation method reveals gene-gene interactions in single-cell RNA-sequencing data

**DOI:** 10.1101/111591

**Authors:** David van Dijk, Juozas Nainys, Roshan Sharma, Pooja Kaithail, Ambrose J. Carr, Kevin R. Moon, Linas Mazutis, Guy Wolf, Smita Krishnaswamy, Dana Pe'er

## Abstract

Single-cell RNA-sequencing is fast becoming a major technology that is revolutionizing biological discovery in fields such as development, immunology and cancer. The ability to simultaneously measure thousands of genes at single cell resolution allows, among other prospects, for the possibility of learning gene regulatory networks at large scales. However, scRNA-seq technologies suffer from many sources of significant technical noise, the most prominent of which is ‘dropout’ due to inefficient mRNA capture. This results in data that has a high degree of sparsity, with typically only ~10% non-zero values. To address this, we developed *MAGIC (Markov Affinity-based Graph Imputation of Cells),* a method for imputing missing values, and restoring the structure of the data. After MAGIC, we find that two- and three-dimensional gene interactions are restored and that MAGIC is able to impute complex and non-linear shapes of interactions. MAGIC also retains cluster structure, enhances cluster-specific gene interactions and restores trajectories, as demonstrated in mouse retinal bipolar cells, hematopoiesis, and our newly generated epithelial-to-mesenchymal transition dataset.

## INTRODUCTION

Single cell RNA-seq (scRNA-seq) is a powerful technology that promises to transform biomedical research. Enabled by recent high-throughput technologies including Dropseq(*1*), Indrop(*2*) and the commercial device 10X chromium3, this data-type is accumulating at a staggering rate. The data offers the promise of addressing a large variety of questions in biology and has successfully been used across many domains to identify novel subsets (*3*-*5*), developmental trajectories (*6*-*8*) and regulatory mechanisms (*9*, *10*).

This data potentially enables the learning of gene-gene relationships in a system-wide scale, based on the naturally occurring variation (*10*-*14*). However, a key challenge underlying the analysis of scRNA-seq is that the observed expression counts capture a small fraction (typically 5-15%) of the transcriptome of each cell, an issue commonly termed “drop-out”(*15*). Such drop-out obscures all but the strongest relationships between highly expressed genes. Thus, while scRNA-seq measures “genome-wide”, the distribution of an individual gene is dominated by measurement noise, which is further exacerbated in lowly expressed genes. However, it is precisely these lowly expressed genes that are often of great interest e.g.: transcription factors and cell surface markers. Researchers have attempted to overcome these challenges by clustering and combining cells, and thus losing the single cell nature of scRNA-seq.

To address these issues we developed MAGIC (Markov Affinity-based Graph Imputation of Cells), a computational approach for recovering gene expression lost due to drop-out. An empowering feature of scRNA-seq data is that while each cell is only sparsely sampled, typical datasets now include thousands of cells and more recently, tens of thousands (*1*, *2*). MAGIC leverages this large sample size to share information across similar cells and fills in missing transcripts by imputing gene expression for any given cell, taking advantage of the rich structure imposed by biology. The resultant imputed data after MAGIC has restored values for likely gene expression in each cell.

We validate MAGIC on bulk and synthetic datasets where the “ground truth” is known. We find that MAGIC restores genes, cells and gene-gene correlations successfully. Further we show that MAGIC preserves cluster structure while enhancing cluster-specific low expression genes in single-cell RNA sequencing data of bone marrow (*16*) and retinal bipolar cells (*3*). We show that in a bone marrow dataset (*16*) collected using MARS-seq2 we can correctly recover the hematopoietic development and the underlying gene-gene relationships. Next, we use in-drop (*2*) to generate new data measuring a breast cancer cell line (*17*) undergoing TGF *β*-induced epithelial-to-mesenchymal transition (EMT). MAGIC reveals a continuous EMT progression with epithelial, mesenchymal, apoptotic, and transitional cells that display a combination of epithelial, mesenchymal and stem-like characteristics. Finally we show that when MAGIC is applied to retinal bipolar cells (*3*) collected using drop-seq (*1*), gene-gene relations are enhanced, while distinct cell sub-types are preserved. This demonstrates that MAGIC can be applied to a wide range of biological systems and single cell technologies.

## RESULTS

Single cell RNA-sequencing is a powerful technology that is marred with the technical challenge of “drop-out”(*15*), where we only observe a small fraction (typically 5-15%) of the transcriptome for each cell (Figure 1A-B). To further complicate matters, this fraction varies greatly between cells due to varying efficiencies of multiple technical steps, including cell lysis, reverse-transcription and subsequent amplification steps. While cell type identification has been relatively successful in face of this sparsity (*1*, *18*), even basic population structure and clustering can be impacted. The impact of drop-out is even more acute when studying gene-gene relationships. Drop-out obscures all but the strongest relationships between highly expressed genes (Figure 1B). To address these issues we develop MAGIC (Markov Affinity-based Graph Imputation of Cells), which receives as input an observed count matrix *D* and shares information across cells to generate an imputed count matrix *D_imputed_*.

**Figure 1:**
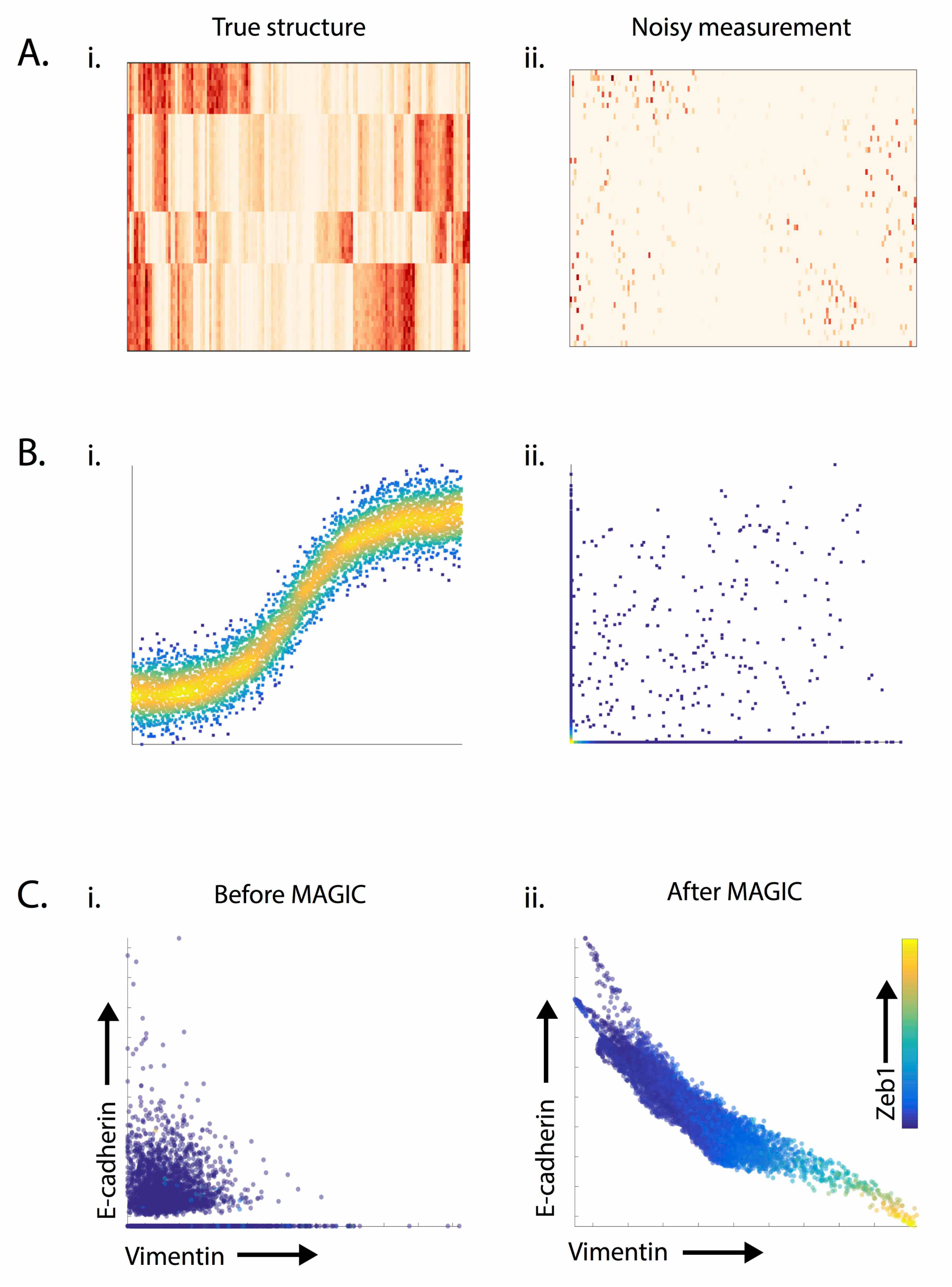
The effects of noise and dropout on the structure of single-cell RNA-sequencing data. A) Synthetically generated gene expression matrix shows the effect of dropout. The matrix in (Ai) demonstrates cluster-structure that is lost in the matrix in (Aii) following “artificial dropout” generated by randomly setting 80% of values to 0. B) Synthetically generated sigmoidal shape of the interaction shown in (Bi) is obscured following drop-out. (Bii) shows the same data following dropout induced by subsampling from an exponential distribution such that 80% of values are set to 0. C) Shows the effectiveness of MAGIC in restoring structure to the canonical EMT gene-gene interaction between Ecadherin and Vimentin from single-cell RNA-sequencing data measuring 7523 HMLE cells 8 and 10 days after TGFB stimulation.

A key property of single-cell data that can be used to alleviate these challenges is that while the number of genes is high, the data is inherently low dimensional due to strong interdependencies between genes(*19*, *20*). First, genes are regulated through shared regulatory programs creating modules of coordinately expressed genes (*20*, *21*), resulting in low rank expression matrices. Moreover, cells are attracted to stable cell-states, driven by feedback loops and other regulatory mechanisms that further constrain possible cell states (*22*-*24*). We find that cell phenotypes are typically limited to dense but smoothly varying “patches”, restricted by biological constraints. The cellular state space thus forms a smooth low-dimensional structure called a *manifold*(*25*). Therefore, while the data is observed in a high dimensional *measurement space,* cell phenotypes can be modeled as residing on a substantially lower dimensional manifold embedded within the measurement space.

We have previously shown that this manifold can be effectively represented using a nearest neighbor graph, where each node represents a cell and edges connect each cell to a neighborhood of its most similar cells, based on their expression. Nearest neighbor graphs have been used to faithfully recover subpopulation structure (*3*, *23*) developmental trajectories (*6*, *7*, *26*) and bifurcations (*7*, *8*). More recently diffusion maps have also been used to find trajectories in single-cell data (*7*, *8*). However, unlike previous work, in MAGIC, we use the diffusion process itself, and its associated *diffusion operator*, to simultaneously discover the manifold and impute gene expression using the manifold structure. Thus by learning the underlying manifold via *diffusion through the data*(*27*), we can restore cellular phenotypes back to the manifold and in the process restore missing transcripts.

The concept of data diffusion is to share information between cells through local neighbors in a process that is mathematically akin to diffusing heat through the data. Data diffusion is implemented via a *diffusion operator* or a Markov-normalized affinity matrix that defines the steps of a random walk through the data. Effectively, this process uses similarities between data points to build a weighted graph structure through which information flows, simultaneously learning the manifold structure. MAGIC uses the graph structure to impute the expression profile of each cell based on the weighted average of its neighbors, smoothing the data and restoring it to its underlying manifold. It is important to note that while we average data across cells, each individual cell retains a unique neighborhood, resulting in a unique expression vector. In Figure 1C, we see that MAGIC reveals the known inverse relationship between Ecadherin and Vimentin during the course of EMT, which is obscured in the original scRNA-seq data due to a high degree of drop-out.

### Overview of the MAGIC Algorithm

MAGIC begins with an *n*-by-m count matrix *D*, representing the observed transcript counts of *n* genes in *m* cells and returns an imputed count matrix *D_imputed_*. The expression of each individual cell, a row in *D*, defines a point in the high-dimensional *measurement space* representing the cell's observed *phenotype.* The counts in the imputed data matrix *D_imputed_* represent the likely expression vectors (phenotypes) for each individual cell, based on data diffusion between similar cells. Key to the success of our graph-based method is to derive a faithful *neighborhood* of similar cells, based on a good similarity metric. We will first describe an outline for MAGIC (see Figure 2 and pseudo-code 1), provide some real world examples of its use and then elaborate on critical details and parameters in later sections.

**Figure 2:**
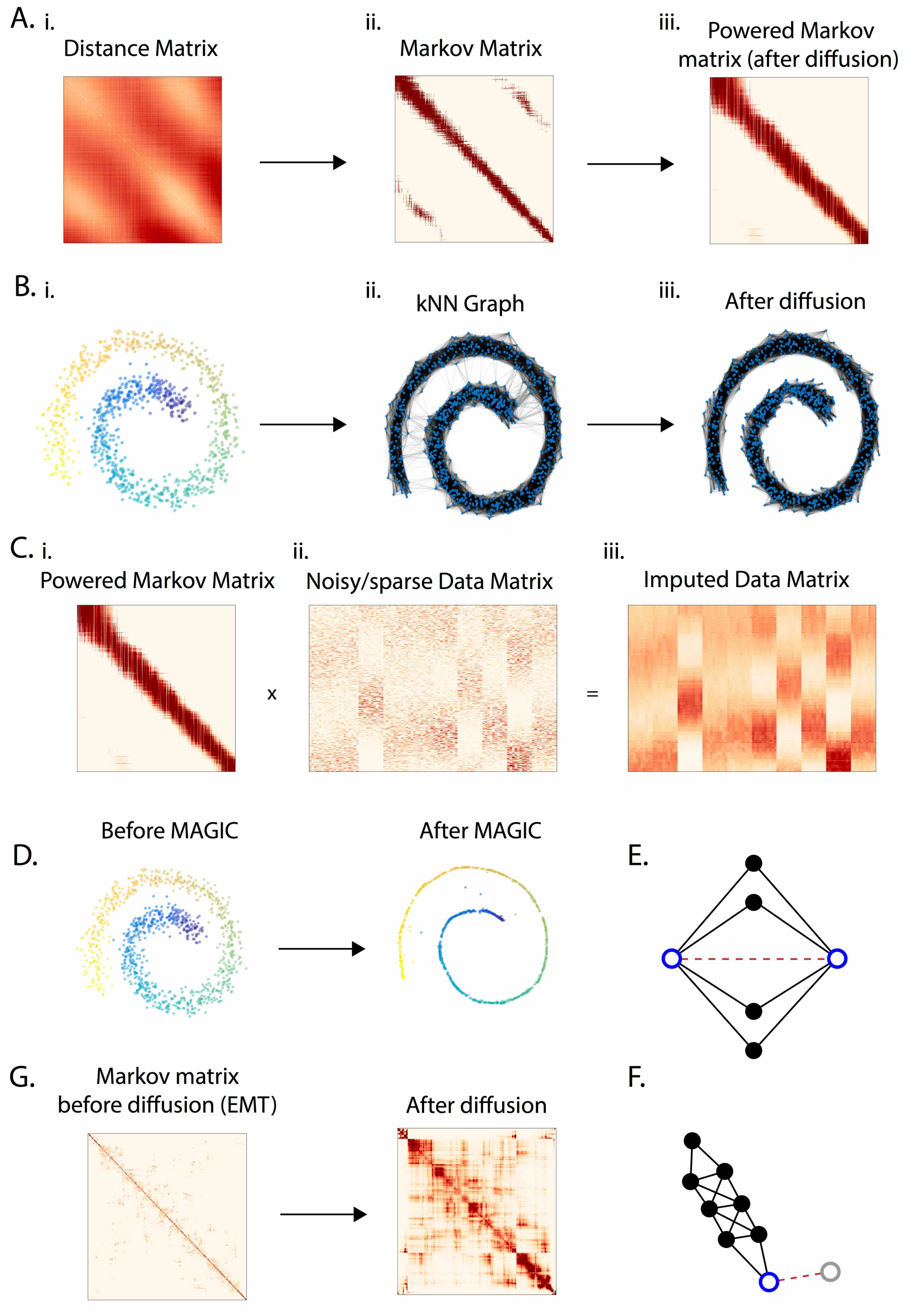
Illustration of the MAGIC algorithm. A) Matrices representing an artificially generated Swiss-roll dataset containing 1000 datapoints. (Ai) A cell-cell distance matrix colored such that brown represents high values and white low values. (Aii) A Markov-normalized affinity matrix computed from the distance matrix via an adaptive Gaussian kernel followed by row-normalization. (Aiii) The matrix in Aii, after it is powered t=16 times. The cells in each of the matrices Ai, Aii, Aii are ordered linearly according to their location on the Swiss roll. B) The synthetically generated Swiss roll dataset is shown in (Bi) as a scatter plot, (Bii) A graph whose edges correspond to the affinities represented in Aii, only nearest neighbors of highest affinity are connected. (Biii) A graph with neighbors determined by the powered Markov affinity matrix shown in (Aiii). C) The imputation step of MAGIC: the diffused Markov affinity matrix (Ci) is right-multiplied by the noisy data matrix (Cii) to obtain the imputed data matrix (Ciii). D) The Swiss Roll dataset before and after MAGIC. E) A toy example where powering the Markov affinity matrix strengthens the connection between two non-proximal neighbors because of the many paths between them. F) A toy example where the powering of the Markov affinity matrix weakens a proximal neighbor affinity because of lack of paths through the region. G) Cluster-structure appears in the powered Markov affinity matrix representing the EMT data shown in 1C.

~~~
 *MAGICS*(*D*, *t*)
     *D* = *preprocess*(*D*)
     *Dist* = *compute_distance_matrix*(*D*)
     *A* = *compute_affinity_matrix(Dist)*
     *M* = *compute_markov_affinity_matrix*(*A*)
     *D_imputed_* = *M^t^* * *D*
     *D_rescaled_* = *Rescale* (*D_imputed_*)
     *D_imputed_* = *D_rescaled_*
 *END*
~~~

A common way to compute similarities between vectors is to: first compute distances (degree of dissimilarity), and then apply a kernel function to obtain similarities from distances. After data processing (in a technology-dependent manner), MAGIC computes a cell-cell distance matrix *Dist* based on Euclidian distance (Figure 2Ai). Distances are then converted into an affinity matrix *A* using a kernel function that emphasizes close similarities between cells. MAGIC uses an adaptive Gaussian kernel, so that similarity between two cells decreases exponentially with their distance. Specifically distances below *σ* (the standard deviation of the Gaussian kernel) are converted into relatively high affinities, which drop off above *σ.* We treat *A* as a weighted adjacency matrix representing the neighbor graph structure where the entry *A(i, j)* represents the edge weight between cells *i* and *j.* We create a Markov transition matrix *M* by row normalizing *A* such that each row sum is 1, representing the probability distribution of transitioning from a particular cell to every other cell in the data in a single step (Figure 2Aii). Raising *M* to the power *t* results in a matrix where each entry represents the accumulated probability of all paths of length *t* between two particular cells, that is *M*^*t*^(*i,j*) represents the probability that a random walk of length *t* starting at cell *i* will reach cell *j* (Figure 2Aiii). Thus we call *t* the "diffusion time”.

Figure 2A illustrates the distance and Markov affinity matrices and Figure 2B the corresponding neighbor graph for a synthetically generated *Swiss roll* with Gaussian noise. Note that while most nearest-neighbor edges follow the spiral, there are *short cut* edges that cut across the spiral. Diffusion maps (*27*, *28*) provide powerful tools that infer the data manifold (a lower dimensional object which embeds the data) based on random walks along the graph's edges. The *magic* in this process is that through the powering of *M*, the underlying structure of the data is revealed. The graph defines small local steps that are limited to very similar cells and the random walk aggregates these small steps into more global intrinsic similarities that follow the data manifold structure. Figures 2Aiii, Biii illustrate the effects of powering the matrix corresponding to the Swiss Roll, spurious edges that cut across the Swiss roll are removed after the matrix is powered, resulting in a graph structure that faithfully follows the Swiss roll.

The imputation step of MAGIC involves learning from the values in neighborhoods resulting from the powered Markov Affinity matrix and right-multiplying *M^t^* by the original data matrix. *D_imputed_ =M^t^* D* (Figure 2C). Technically, when a matrix is applied to the right of the Markov Affinity matrix it is considered a *backward diffusion operator* and has the effect of replacing each entry *D(i, j)* that is gene *j* in cell *i,* with the weighted average of the values of the same gene in other cells (weighted by *M^t^*).

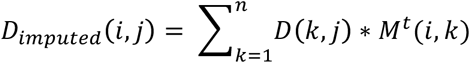

The weighting is done precisely by the affinity *M*(*i*, *k*) of cell *k* to cell *i*. Since each row of M sums to 1, the values of *D_imputed_*(*i*,*j*) do not increase arbitrarily. This process effectively restores the missing data to reside on to the underlying manifold structure, which embeds the majority of the data. In Figure 2D we see the Swiss roll data before and after imputation. Following imputation, the data points almost perfectly reside within the 1-dimensional Swiss roll used to generate the original data matrix.

While the powered Markov affinity matrix increases the number of cell neighbors, reweighting also occurs: dense areas of the data result in more possible paths and thus weights are concentrated in these areas (Fig 2E-F). Thus the affinity between the two blue points highlighted in 2D is increased by diffusion, due to many common paths. Similarly the affinity between the blue and gray points in Figure 2F is downweighted due to a lack of paths in this region. Therefore, paths through dense areas in the data increase their weight, and paths through sparser areas (often spurious edges that result from noise) are weighed down. Such effects are not limited to toy examples. In Figure 2G, we illustrate the strong cluster structure that emerges in the powered Markov matrix representing 1500 cells undergoing EMT.

In the final step of MAGIC, we re-scale the count matrix. The MAGIC process resembles heat diffusion in the graph, which has the effect of spreading out molecules, but keeping the total sum constant. This in effect means that the average value of each molecule decreases after imputation. To match the observed expression levels (per cell), we rescale the values so that the max value for each gene equals the 99^th^ percentile of the original data. Thus those cells with high expression of a gene are brought up to similar levels as the original data and all other values are proportionally scaled up with them.

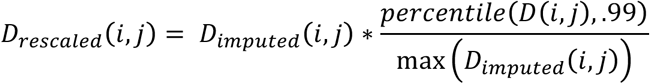

### MAGIC Enhances Cluster and Trajectory Structures in Bone Marrow

To evaluate MAGIC in a real data setting we applied it to a recently published scRNA-seq mouse bone marrow dataset(*16*), collected using the plate based MARS-seq2 protocol (*18*). Bone marrow is a complex tissue containing a variety of well-studied cell subtypes undergoing differentiation and is therefore an ideal test case. In (*16*), the authors identify 19 clusters (shown in Figure 3A-B): mature erythrocytes (cluster 1), precursor cell types such as early erythrocytes, and early monocytes (clusters 7-12), more mature myeloid cell types such as monocytes (cluster 14, 15), and neutrophils (clusters 16, 17). While clustering worked reasonably well, the data matrix is quite sparse and most cells are missing many canonical genes, known to be expressed in their respective cell types.

**Figure 3:**
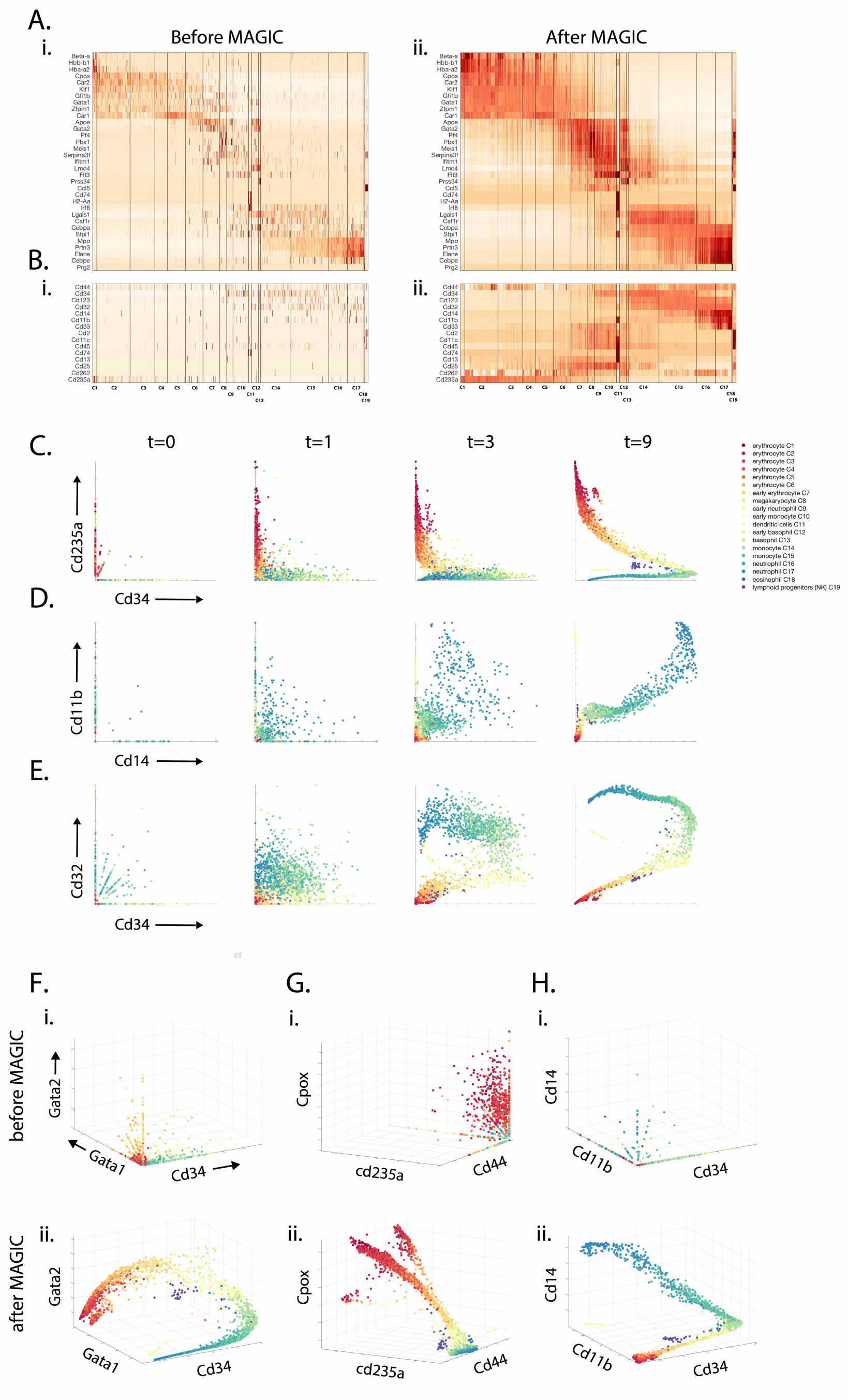
MAGIC applied to single-cell mouse myeloid progenitor data: We use the mouse bone marrow dataset from (*16*)). A) The gene expression matrix for key hematopoietic genes (as identified by (*16*)) before MAGIC (Ai) and after MAGIC (Aii). Genes are visually grouped by hierarchical clustering. MAGIC shows enhanced expression of cluster specific markers while still maintaining cluster structure. B) The gene expression matrix for characteristic surface markers of immune cells, before (Bi) and after (Bii) MAGIC. C) Scatter plots demonstrating the recovery of the Cd34 vs Cd235 relationship after different amounts of diffusion (*t*=0,1,3,9). In these scatter plots, dot represents a single cell, plotted according to its expression values (measured at *t*=0 and imputed for *t*=1,3,9), colored based on the clusters identified in (*16*)). After imputation, clear gene-gene relations emerge in the data and the clusters form a continuous progression. Only the erythroid lineage cells (orange-red) increase in Cd235 with the decrease in Cd32. D) Shows the imputation of the relationship between myeloid markers Cd11b and Cd14, which are co-expressed in neutrophils (shown in blue) and later myeloid cell types. E) Shows the imputation of the Cd32 vs Cd34 relationship, which has diverging behavior for the erythroid lineage (shown in warm colors) as compared to the myeloid lineage (shown in cool colors). F-H) Show before and after MAGIC of 3-D relationships (with diffusion time *t*=9).

Figure 3Ai,Bi shows the expression levels of characteristic genes from these clusters before and after MAGIC. While these genes have relatively high expression levels when detected, their expression is spotty and inconsistent within their respective clusters (Figure 3Ai). For instance, Hba-a2 (the alpha unit of hemoglobin) and Hbb-b1 (beta unit of hemoglobin) are only expressed in erythrocyte clusters (C1-C6) in 36% and 17% of cells. Cebpa, a neutrophil transcription factor is only expressed in 52% of neutrophil clusters (C17-17). We applied MAGIC to this data (with parameters *npca=100, ka=4, t=9,* Figure 3B). After MAGIC we see more consistent expression patterns, all cells significantly express hemoglobin subunits and Cebpa in their respective clusters.

Figure 3Aii,Bi shows the expression of canonical surface markers, before and after MAGIC. These markers are typically used to identify and gate these immune populations at the protein level. However, at the transcript level they are typically lowly expressed (likely because these are relatively stable proteins). Without knowing the clustering associations, MAGIC is able restore many of the proper surface markers to each cell. Before imputation, the monocyte clusters C14, C15 have only 1.6% cells expressing Cd14 and 5.8% cells expressing Cd11b and only 10% of the dendritic cells (cluster C11) express Cd32. After we apply MAGIC, 94% of monocytes express Cd14, 98% express Cd11b and 97% of dendritic cells express Cd32. Thus we see that MAGIC preserves the cluster structure and enhances the expression of cluster-specific markers.

The sparsity of the data is even more evident when viewing the data using the biaxial plots typically used to analyze immune subsets. Prior to imputation, most of the data lies on the axes (Fig 3C-E, *t*=0). Given the degree of dropout, it is rare for both genes to be observed in any given cell, thus obscuring relationships *between* genes. MAGIC restores these missing values and relationships, recreating the biaxial plots one typically sees in flow cytometry. Fig. 3C shows that as cells mature from early progenitors into erythrocytes, Cd235b (an erythrocyte marker) increases and they gradually lose Cd34 (a hematopoietic marker). Fig. 3D shows the positive correlation between Cd14 and Cd11b, which are both upregulated as monocytes mature, a relationship that is undetectable in the raw data. Fig 3E shows the inverse relationship between Cd32 (Fcgr2b) which is an inhibitory receptor expressed on mature dendritic cells and the hematopoietic marker Cd34 for the myeloid/dendritic branch, and positive correlation between the two in the erythroid branch.

By superimposing the reported clusters onto the biaxial plots we see that cells in the same clusters are grouped closely together and gene-gene relations gradually change between clusters as the cells mature and differentiate. MAGIC not only maintains the cluster structure; it is also able to recover finer structural details. Figures 3C-E also demonstrate the effects of powering the Markov matrix. After a single step (*t*=1) some imputation occurs, but there is hardly structure in the data. At *t*=3 a shape reminiscent of the final imputation begins to emerge. However, it is quite noisy. At *t*=9, the optimal parameter chosen by our correlation dimension based method, the structure is clear and well formed.

Figures 3F-H show gene-gene relationships in 3 dimensions. While little structure is visible before MAGIC, as in 2D, after MAGIC we observe the emergence of a continuous developmental organization that shows connections between clusters. Gata1, a TF essential for erythrocyte and megakaryocyte lineage commitment (*16*), shows gradual increase for clusters 1-8 as Cd34 decreases (Figure 3Fi,ii). Similarly, early progenitor TF Gata2 goes up with erythrocyte development. However, as erythrocytes reach maturity, Gata2 is repressed. Cd235a (an established erythroid marker) and Cpox (an enzyme associated with hemoglobin production), both increase with erythroid development (Figure 3Gi,ii). Heterogeneity in erythrocytes is highlighted by Cd44, which is known to be variably expressed in erythrocytes dependent on age of a cell(*29*). Figure Hi,ii shows characteristic myeloid markers Cd14 and Cd11b as a function of Cd34. Dendritic cells high in Cd11b. Monocyte and neutrophils high in both Cd14 and Cd11b.

### MAGIC Reveals Progressions in EMT Data

To demonstrate MAGICs ability to reveal fine structure and temporal ordering of events during cell state transitions, we chose to examine the epithelial-to-mesenchymal transition (EMT). We profiled ~7500 HMLE cells (*17*), 8 and 10 days following TGFβ induction of EMT, collected using the in-drop microfluidic protocol (*2*) (see Experimental Methods). Induction of EMT is asynchronous and each cell progresses along the transition at a different rate. Therefore, on days 8 and 10, cells occupy all phases of EMT.

Even the most canonical relationship, the decrease in E-cadherin (epithelial marker), coinciding with an increase of Vimentin (mesenchymal marker), is obscured due to dropout. After MAGIC is applied to this data (with parameters *npca=20, ka=10, t*=6) the relationship between E-cadherin and Vimentin is successfully recovered (Figure 1C). The formation of structure becomes even more pronounced when looking at the data with a 3^rd^ canonical marker, Fibronectin (mesenchymal marker). Before MAGIC, most cells lie near the origin of this 3-dimensional plot (Fig. 4A) and after MAGIC structure emerges (Fig. 4B). We can interpret this structure by overlaying the expression of specific genes onto the cell scatter-plot. For example, expression of Zeb1, a key transcription factor known to induce EMT (*30*), progressively increases as Vimentin and Fibronectin increases and the cells become more mesenchymal.

**Figure 4:**
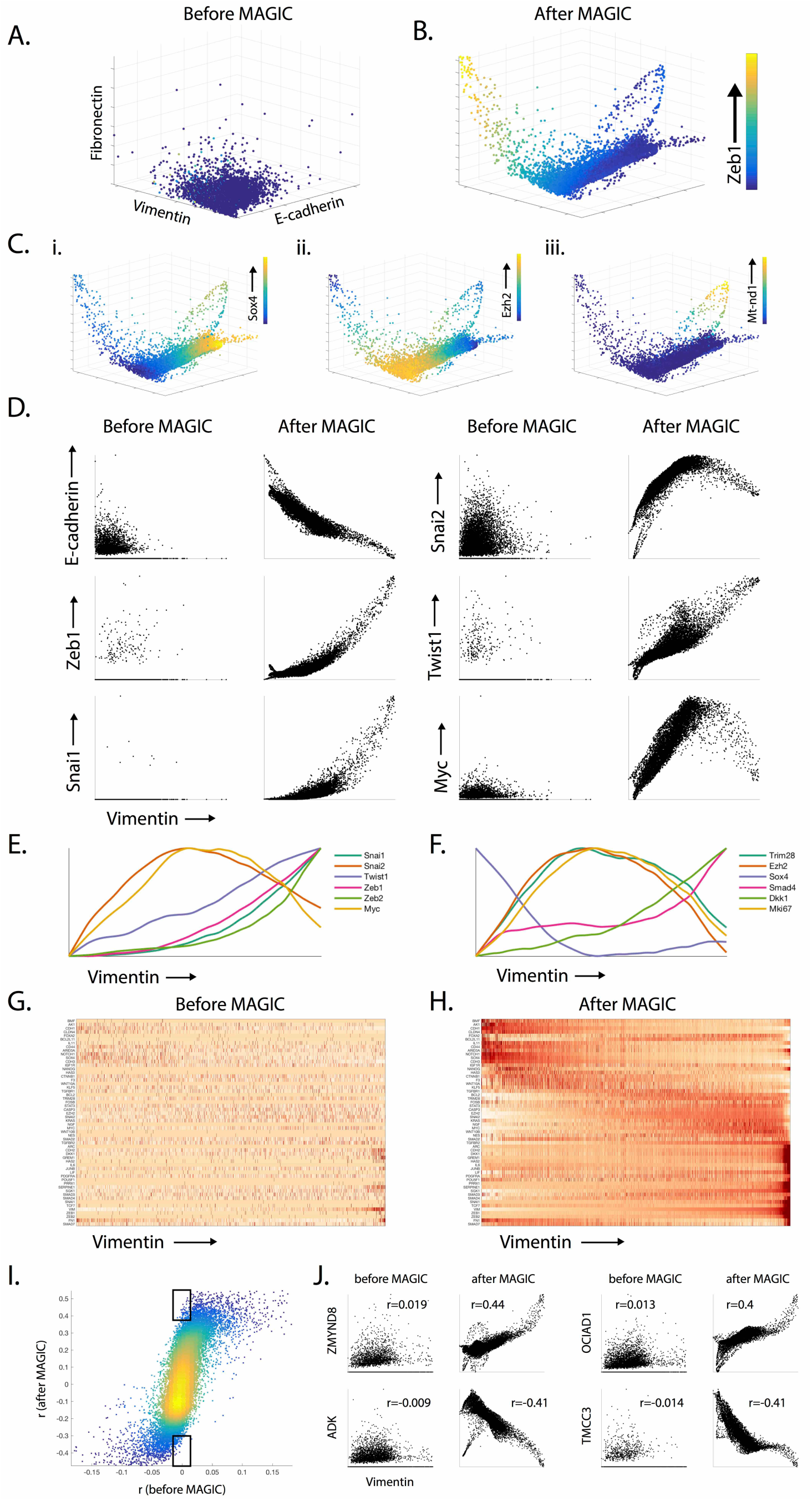
MAGIC recovers gene-gene interactions in epithelial-to-mesenchymal transition data. This data was collected on Indrop 8-10 days after TGFB-stimulation of HMLE breast cancer cells (see Methods). A-B) 3-D scatterplots between canonical EMT genes Ecadherin, Vimentin, and Fibronectin. A) Scatterplot before MAGIC with most cells near the origin, B) After MAGIC, the scatterplot reveals a continuous trajectory. The cells are colored by the level of Zeb1, a key EMT transcription factor, which increases through the middle of the mesenchymal branch. C) The Ecadherin-Vimentin-Fibronectin relationship, colored by other genes: Ci) Sox4. Cii) Ezh2, a chromatin modifier known to be involved in EMT Ciii) mt-nd1, a mitochondrial gene which is only accessible if cells are undergoing apoptosis involving MOMP (mitochondrial outer membrane permeabilization), D) Scatter plots of several well known EMT genes against Vimentin, before and after MAGIC. E-F) Temporal trends of gene expression using Vimentin as a proxy for EMT progression. (E) Lowess regression plot (with window size 150) showing temporal trends of transcription factors associated with EMT. (F) Lowess regression plot of additional genes as indicated. G-H) Gene expression matrix of Hallmark EMT genes shown as a heatmap. H) Gene expression matrix with cells ordered by vimentin post-MAGIC. I) Gene-gene correlation (r) before MAGIC vs gene-gene correlation after MAGIC. The range of (r) before MAGIC is roughly [-0.15, 0.15]. Range increases to [-0.4, 0.5] after MAGIC. A large number of gene-gene interaction show increased correlation after MAGIC. Top black box shows a subset of such genes. Bottom black box shows genes that remain with low correlation after MAGIC. J) Genes Zmynd8 and Ociad1 acquire significantly increased positive correlation with Vimentin after MAGIC and genes Adk and Tmcc3 that acquire significant negative correlation with Vimentin after MAGIC.

Figure 4C shows several other prominent genes that mark different cell states, displaying coherent expression patterns that help interpret the different regions of the plot. The epithelial state (high E-cadherin) is marked by high expression of Sox4 (Figure 4Ci), which has been shown to bind to the promoter of Ezh2(*31*). Looking at the expression pattern of Ezh2, we see that it is high in an intermediate state (Figure 4Cii), after the cells begin to transition and lose E-cadherin, but before they gain Vimentin. Thus, the temporal ordering of events between expression of Sox4 and the rise of Ezh2 is restored by MAGIC. A particularly fine structure revealed by MAGIC are two branches that deviate from the main trajectory, which display an increase in mitochondrial DNA, reflecting a progression into apoptosis (Figure 4Ciii). Since mitochondrial transcripts are generally inaccessible to the mRNA capture protocol, their presence indicates that the cells have undergone MOMP (mitochondrial outer membrane permeabilization)-a key step in apoptosis. The apoptotic state is supported by the rise of additional apoptotic markers in these cells including Cox5bp6 and Tnf (data not shown).

TGF*β* is known to upregulate mesenchymal transcription factors like Twist, Zeb, and Snail (*30*). Due to low expression of transcription factors, the raw data does not allow for quantification of relationships between genes and their progression along EMT, as is apparent from Figure 4D (before MAGIC). However, following MAGIC relationships between known EMT genes are revealed, recapitulating their known temporal trends along EMT (we use the mesenchymal gene Vimentin as a proxy for progression, due to its monotonic increase during this process). We observe that Zeb1, Snai1, and Twist1 increase as Vimentin goes up. Cdh1 (E-cadherin) goes down. Myc and Snai2 increase and subsequently decrease as cells become mesenchymal (Figure 4D). We see that Snail2 rises first and peaks at an intermediate state, followed by Snail1 and Zeb1 which rise along with Vimentin at later stages of the transition (Figure 4D).

The fine resolution of the inferred data allows for the investigation of temporal ordering in gene expression dynamics along EMT. We removed apoptotic cells and ordered the remaining cells based on their imputed Vimentin levels and plotted the expression levels for known EMT regulated genes (Fig. 4 E-F, see methods). This illustrates the diverse response associated with EMT, in which expression of genes such as Ezh2, Trim28 and Myc precedes up-regulation of Twist1, Zeb1 and Smad4.

The single cell and genome wide resolution combined with MAGIC enable a systemwide investigation of the dynamics of gene expression during a state transition, in our case along EMT progression, as shown in the heatmaps of Figure 4G. In Figure 4G, genes are ordered by when these peak along EMT after imputation (Figure 4H), showing three general classes of genes, one that is high in the epithelial state (e.g. Cdh1), one that is high in an intermediate state (e.g. Trim28), and one that is high for the mesenchymal state (e.g. Dkk1).

For the majority of gene-gene relationships that we investigated the pre-MAGIC bi-axial plot is highly non predictive of the post MAGIC trend, e.g. whether the genes are positively or negatively related. To quantify whether the original data is predictive of the trends that emerge after imputation, we quantified Pearson correlation between Vimentin and all other genes, before and after MAGIC (Figure 4I). First we observe that the range of correlations is much lower before MAGIC, −0.15 to 0.15 versus −0.5 to 0.5 after MAGIC. Moreover, we find that if the correlation is (relatively) high before MAGIC the correlation will also be high after MAGIC, i.e. MAGIC does not remove preexisting correlations. However, most relationships have relatively low correlation before MAGIC but for a large number of these the post MAGIC correlation is high. These relationships would not have been detected in the original data. The boxes in Figure 4I mark regions of low (-0.02 to 0.02) correlation before MAGIC and high (>0.4 or <−0.3) correlation after MAGIC. Figure J shows 4 relationships picked from those regions where before MAGIC correlation is low but after MAGIC correlation is high.

### MAGIC Preserves Cluster Structure in Retinal Bipolar Cells

We have shown that MAGIC enhances trajectory structure both in the case of hematopoiesis (Figure 3) and the EMT cell state transition (Figure 4). To demonstrate MAGIC's ability to impute given distinct and separated cell types, we use a mouse retina dataset collected with the drop-seq protocol (*3*), the third scRNA-seq technology on which we demonstrate MAGIC.

The authors of (*3*) investigate phenotypically distinct bipolar cell types by means of several clustering algorithms. Of these methods they concluded that Jaccard-Louvain (also known as Phenograph(*23*)) performed best. Therefore, we used this approach and clustered the cells (based on the original count matrix) with Phenograph (*k*=30). Then we ran MAGIC (*npca*=100, *ka*=10, *t*=6) and visualized pre-MAGIC cluster structure on the imputed data. Figure 5A shows gene expression of a set of key genes (as identified by the authors) ordered by Phenograph clusters. Because this data has a clear cluster structure, we wanted to verify that MAGIC does not artificially connect clusters. To quantify this we re-clustered the after MAGIC data (Phenograph, *k*=30) and computed the Rand index (measure of similarity between clustering solutions (*32*)) between the before MAGIC and after MAGIC clusters, resulting in a Rand index of 0.93. Rand index has a value between 0 and 1, 1 indicating maximal correspondence.

**Figure 5:**
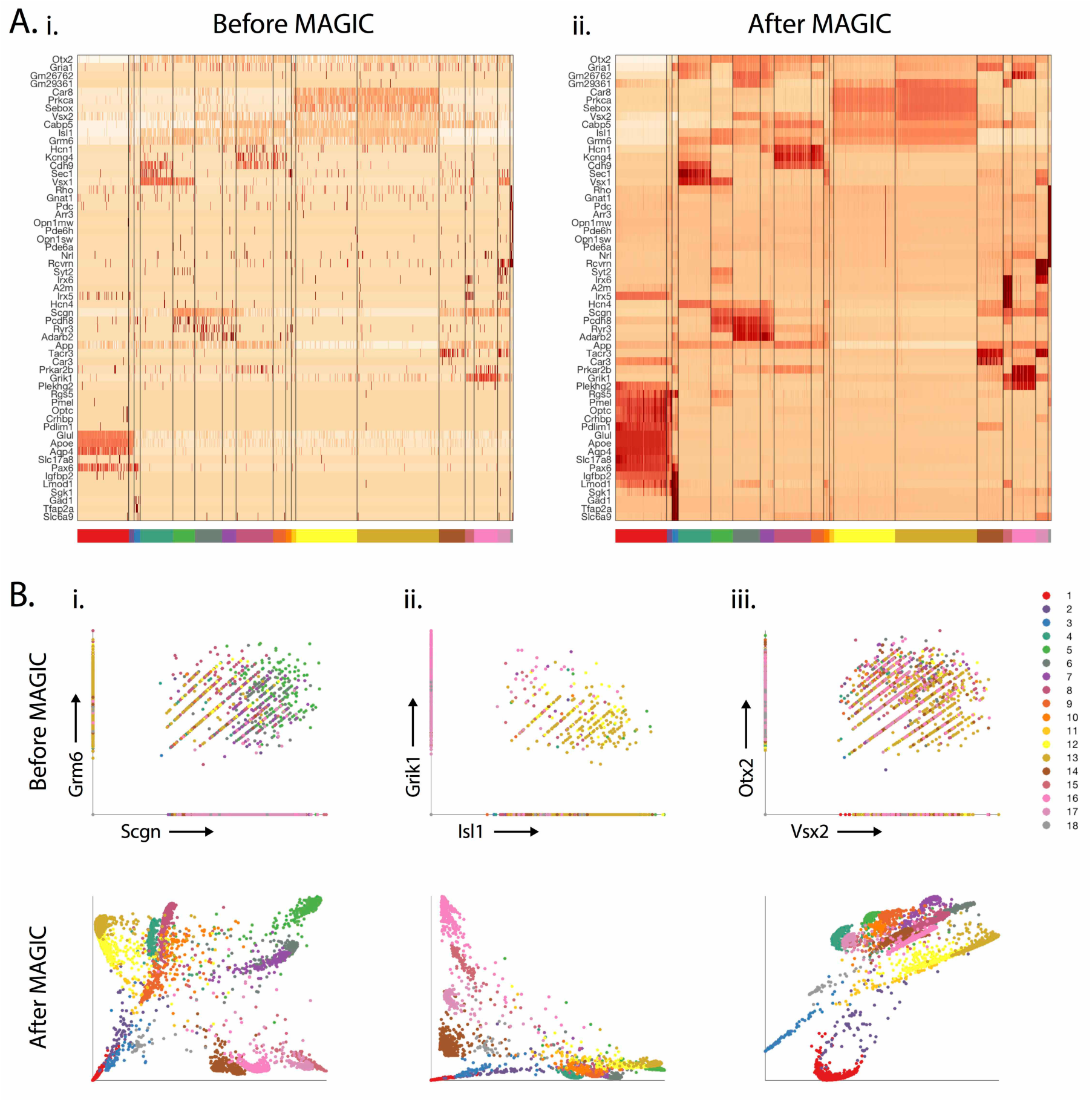
MAGIC applied to mouse retinal bipolar cell data. We use the mouse retina dataset from (*3*). A) Gene expression matrix of 6174 mouse retinal bipolar cells (one replicate of the data, with cells containing <1K transcripts discarded) before imputation. The cells (in columns) are clustered using Phenograph(*23*). Genes (in rows) are ordered using hierarchical clustering. After MAGIC, Cluster-specific genes such as Optc for cluster1 show more consistent expression within the cluster but do not show expression outside the cluster. B) Scatter plots demonstrating several 2D relationships on this data before and after MAGIC. Each cell is colored according to its cluster id. The relationships show differing trends based on cluster id. For instance, the Scgn-Grm6 relationship shows a positive trend most clusters (but with different marker ranges) but negative trend in the brown and pink clusters.

As additional visualization, we plotted various gene-gene interactions before and after MAGIC, and colored cells by their before MAGIC cluster. MAGIC reveals distinct relationships between the genes. However, unlike bone marrow or EMT, the expression levels create distinct and separated regions in the scatter plot corresponding to each of the clusters.

For example, Figure 5Bi shows the relationship between the ON bipolar cone markers Scgn and Grm6. We see that the two markers relate to each other differently in different clusters of cells. For instance, in clusters 5 (green), 6 (gray) and (*7*) purple Scgn and Grm6 are both highly expressed, and moreover within these clusters the two markers show a positive relationship. Clusters 14-17 have high expression of Scgn and low expression of Grm6 and show a negative relationship within the clusters. Figure 5Bii shows that Isl1 and Grik1 are never both highly expressed in the same clusters. Clusters 4-13 are high in Isl1 but low in Grk1, clusters 15-17 are high Grk1 but low in Isl1, clusters 1-3 express lsl1 at low levels but have Grk1 off. Moreover, the level of one or the other marker is distinct for different clusters, and therefore the clusters are separated here. Vsx2 and Otx2, pan-BC markers, are co-expressed in most bipolar cells (Figure 5Biii) to distinct levels. Clusters 4-18 express both markers whereas clusters 1-3 are low in both markers.

Therefore, we can conclude that MAGIC does not artificially connect clusters but enhances gene expression signals within each cluster.

### Constructing MAGIC's Markov Affinity Matrix

Now that we have demonstrated a few examples of how MAGIC enhances signal in scRNA-seq datasets, we will describe the algorithm in more detail. One of the most critical steps in MAGIC is computing the affinity matrix *M. M* defines the graph structure and cell neighborhoods; MAGIC can only succeed if this faithfully represents the geometry of the data. Thus, diffusion-based methods fail or succeed based on how the affinity matrix is constructed. Our construction of *M* is based on the following steps:

1. Computation of a cell-cell distance matrix *Dist*
2. Computation of the affinity matrix *A* based on *Dist*, via an adaptive Gaussian kernel function
3. Symmetrization of *A* using an additive approach
4. Row-stochastic Markov-normalization of *A* (so each row sums to 1) into Markov matrix *M*.

Computation of the distance matrix is straightforward, starting from a pre-processed data matrix *D*, a cell-cell Euclidean distance *Dist* is computed for each pair of rows in *D*. Then, the affinity matrix *A* is computed from *Dist* using a kernel function that converts distances into affinities based on the Gaussian Kernel, as follows:

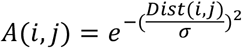

With a negative double exponential function, distances beyond the standard deviation σ rapidly drop off to zero affinity and hence the choice of σ, the *kernel width,* is a key parameter. While MAGIC is relatively robust to the exact value of σ (Fig. 10), this value needs to be within a reasonable range. If σ is too small, the graph become disconnected leading to noise and instability. If σ is too large, distinct and distant phenotypes will be collapsed and averaged together, losing resolution and structure in the data. However, cell phenotypic space is not uniform: a stem cell can be orders of magnitude less frequent than a mature cell type and transitional cell states are also rare. Therefore, σ that would be appropriate for a mature cell type would be far too coarse to capture fine details of the differentiation in the more progenitor cell types.

Additionally, denser phenotypes can dominate the imputation. Cells in dense areas have more neighbors and therefore exert more influence than cells with fewer neighbors. Moreover, dense phenotypes are further reinforced by multiple similar cells with large neighborhoods. During diffusion, dense phenotypes iteratively attract more and more cells towards them and dominate the data. We see such an example in Figure 6, each iteration attracts more and more cells towards the dense center. To preserve the more rare cell states, we need to equalize the *effective* number of neighbors for each cell, thereby diminishing the effect of differences in density. Instead of fixing a single value for the kernel width *σ*, we adapt this value for each cell, based on its local density. Specifically, to equalize the number of neighbors we set the value *σ(i)* for each cell *i* to the distance to its *kath* nearest neighbor:

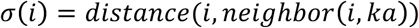

**Figure 6:**
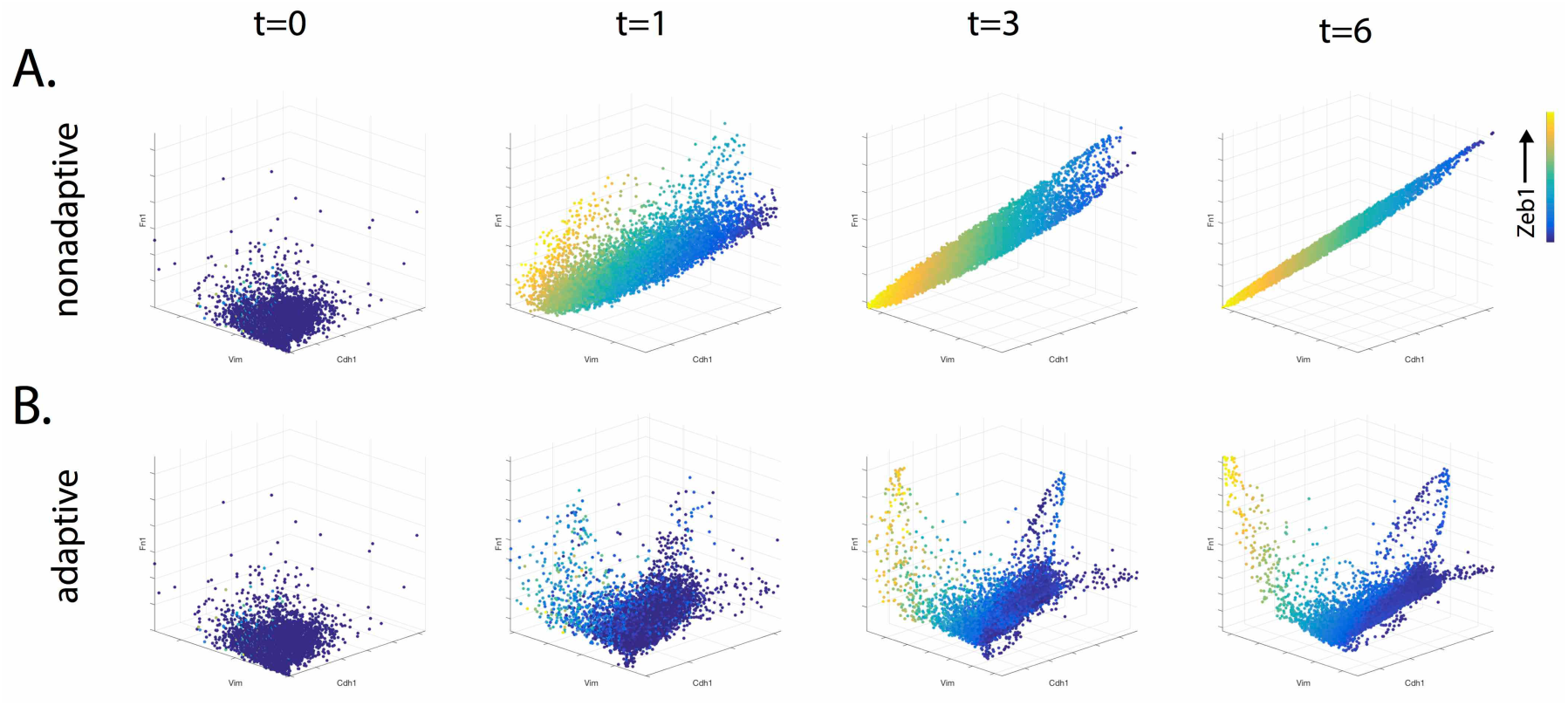
Comparison between adaptive and non-adaptive kernels. A) 3D-scatterplot of Ecadherin, Vimentin and Fibronectin, colored by Zeb1 after MAGIC performed with different amounts of diffusion (*t*=0,1,3,6) using a non-adaptive kernel. The fixed kernel alters and deforms the shape of the data as cells in denser regions have more neighbors, while cells in sparse areas have fewer cells to exchange information. B) The same data with the adaptive kernel used in MAGIC revealing finer structures in the data.

Thus the kernel is wider in sparse areas and smaller in dense areas. To maximize our sensitivity to recover fine structure, we choose *ka* to be as small as possible, such that the graph remains connected. Comparing non-adaptive to the adaptive kernel on the EMT data in Figure 6, we see that the non-adaptive kernel coarsely captures only the single strongest trend in the data, whereas the adaptive kernel does not collapse the data, but rather imputes finer structures. For example, the adaptive kernel identifies the two apoptotic trajectories in the data. Thus, the use of an adaptive kernel is critical for MAGIC's success. To improve computational efficiency and robustness, we ensure sparsity in the resulting affinity matrix A and allow each cell to have at most *k* neighbors. Since the standard deviation of the kernel bandwidth is set locally to the distance to the ka-th neighbor we set *k* = *3ka* to ensure that the *k*NN graph covers the majority of the Gaussian kernel function. All additional affinities (which are already close to zero) are set to zero.

Another important component towards MAGIC's success is the quality of the *diffusion process* that occurs when the affinity matrix is powered. A good process would smooth the data in a manner that follows the shape of the underlying manifold. It has been shown (*33*) that to mimic a discretized diffusion that achieves these properties, the affinity matrix must be symmetric and positive semidefinite, with eigenvalues in the range of zero to one. Negative eigenvalues would simply flip back and forth at each powering, leading to instability. With values greater than one, things would be sensitive to outliers and powering would wildly amplify.

The adaptive kernel results in an asymmetric affinity matrix where *A*(*i*, *j*) *≠ A(j, i),* which we need to symmetrize in order to achieve these desired properties for *A*. We take the additive approach to symmetrization, which averages the affinities, helps pull in outliers and denoises the data. We construct the symmetric affinity matrix as:

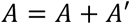

The final step is the row-stochastic normalization that renders the affinity matrix into a Markov transition matrix M. Each row represents a probability distribution, where *M(i,j)* is the probability of cell *i* transitioning to cell *j.* Each row must sum to 1, which we achieve simply by dividing each entry in A by the sum of row affinities.

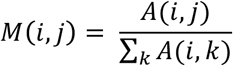

We note that we want a cell's own observed values to have the highest impact on the imputation of its own values, thus our transition matrix allows for self-loops and these are the most probable step in the random walk. The distance between a cell and itself is zero, therefore its weight in the affinity matrix before normalization is 1 (regardless of *σ)* ensuring the measured values in each cell retain a high weight in its imputation.

### Choosing diffusion time for the Markov Affinity Matrix

Another key component of MAGIC is the "diffusion” that occurs when the Markov Affinity Matrix is powered, before the imputation step *D_imputed_ =M^t^ * D.* Since we start with a relatively small *k,* the neighborhood defined by *M* is small and not sufficiently robust. After the powering of *M,* phenotypically similar cells should have a strong weighted entry, whereas spurious neighbors are down-weighted (Figure 2C,D) and therefore diffusion (powering) is particularly well suited to correct for the effects of dropout typical in scRNA-seq.

We assume that the data lies on a lower dimensional manifold, which is obscured by drop-out and additional sources of noise in the data. The true manifold structure of the data is captured by the top eigenvectors of *M,* whereas the rest of the eigenvectors likely represent noise. Powering has the effect of reducing noise in the Markov matrix while retaining biological signal. As Eigenvalues lie in the range [1,0], these are gradually reduced by the powering operation. The number of dimensions encoded in *M^t^* decrease as *t* increases, reducing the noise dimensions to zero and effectively learning the structure of the data manifold. Prior to imputation, with so many values set to zero (due to drop-out), a large fraction of the cells reside well outside of the data manifold, effectively residing as singletons with dimension zero. When *t* is in the right range, powering has the effect of reducing noise and *D_imputed_ =M^t^ * D,* effectively brings noisy and outlier points into the data manifold structure and increases the local density and dimension of most cells. Larger values of *t* begin to impact the manifold related eigenvectors, leading to the decrease in true dimensions of the data. Such values of *t* will result in *over-smoothing* and loss of information after imputation. Therefore, an optimal choice of *t* is a value that recovers the manifold structure, so that cells reside in the full range of this structure.

A good measure for this is *correlation dimension (34),* a measure that estimates the average rate of increase (over all points x) of neighbors within radius *r,* as *r* grows. The main idea behind correlation dimension is that when the intrinsic dimensionality of the data is m, since there are *m* different directions in which close data-points might accumulate, the probability density *p*(*x*, *r*) inside a ball of radius *r* centered at a point x scales as *r^m^* for small *r.* Correlation dimension captures two things, how well does the data reside in distinct densities and the intrinsic dimensionality of these structures. We choose the *t* that will maximize the correlation dimension of the imputed data. Initially, as noise and sparsity dominate the data, the correlation dimension will be low as many points are isolated singletons with dimension 0. Upon initial diffusion and imputation, the noise dimensions are diminished. Therefore, data-points come closer together into the manifold, increasing the neighborhood density and with that the correlation dimension also increases. Beyond an optimal *t*, diffusion begins to reduce the manifold dimensions, decreasing the intrinsic dimensionality of the data and subsequently the correlation dimension.

We search for a *t* that maximizes the correlation dimension. The fraction of points at a radius *r* from a point *x* is denoted *p*(*x*, *r*), and the average of this over all points is denoted *c*(*r*). In a dataset with *N* points, we give the following definitions:

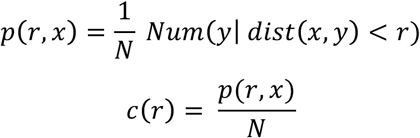

To measure the exponent in the rate of increase in *c*(*r*) vs *r*, we take the slope of the log-log plot of *c*(*r*) vs *r.* We estimate this slope using the following procedure. We first measure the first-nearest neighbor distance for all points and set *r* to the maximum of these distances, so that each point will have at least one neighbor. Then we increase *r* in small step increments to the maximum value, which is the diameter of the data. Thus we probe values in the range [*rmin*, *rmax*] where:

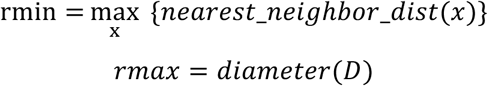

Following this computation, we have a vector *R* of points, for which we compute the slope between *R* and *c(R):*

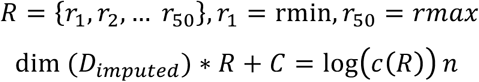

Using this method, we can compute the correlation dimension for a range of *t*, starting at *t =* 1. In Figure 7A, depicting EMT data, we observe that the correlation dimension initially rapidly increases, then peaks, and continually decreases thereafter. We also observe the relationship between *ka* (effective number of neighbors) and *t,* the larger the neighborhood, the quicker the correlation dimension peaks and subsides. In figure 7B we illustrate the resulting imputed data for different combinations of *ka* and *t,* with the peak time-point highlighted in green. The data first becomes less noisy and gains structure, following a peak at *t*=6, after which the data manifold begins to contract. We can also observe that this process does not occur too rapidly and the results are largely similar across a range of values for both *t* and ka. Figure 7C,D shows a similar representation for the bone marrow data, however, in this case the process is imperfect and the imputation seems best at a second peak in correlation dimension. Thus, while correlation dimension acts as a good guide, it is best to look at the data, rather than fully automate the procedure.

**Figure 7:**
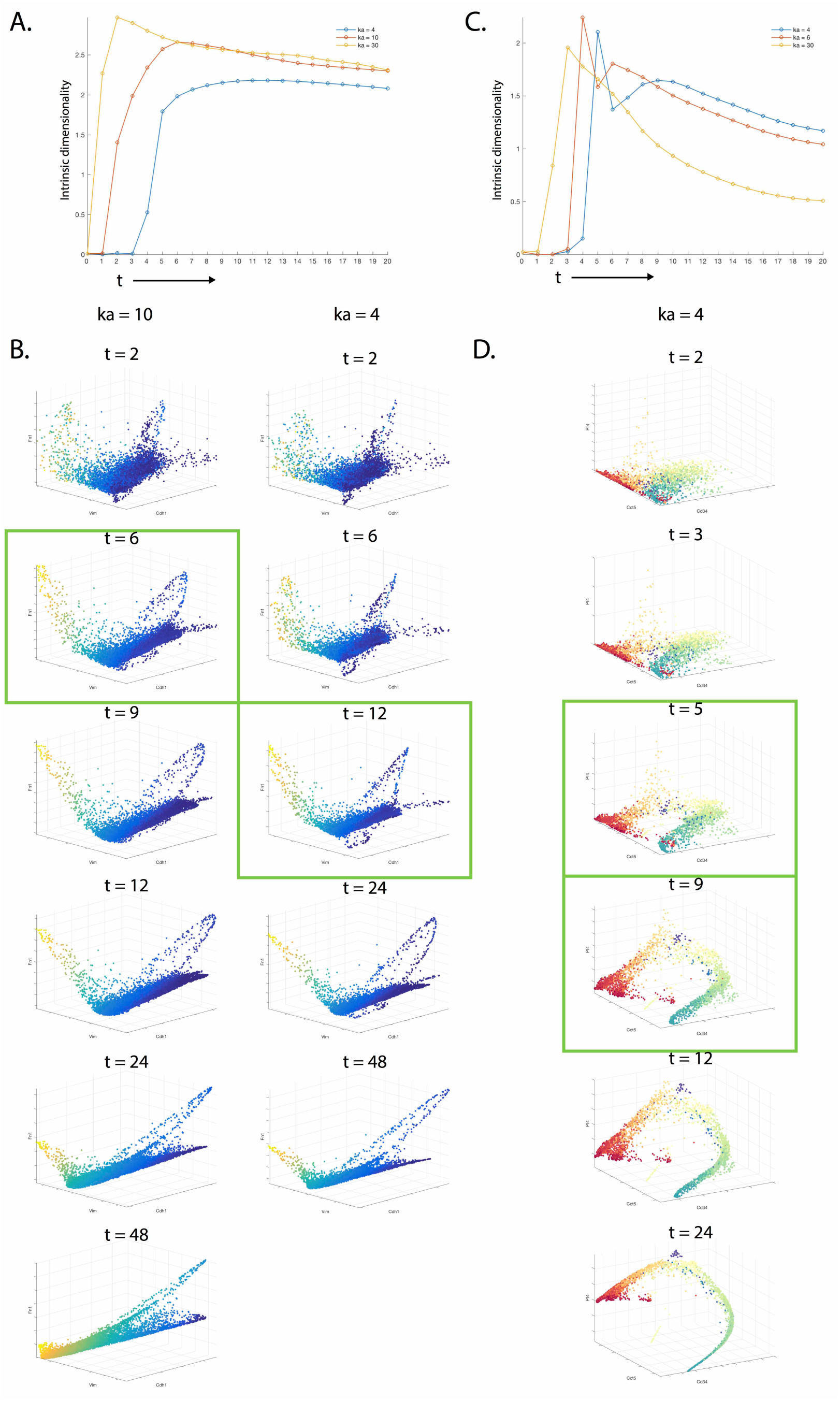
Finding the optimal diffusion time (*t*) using intrinsic dimensionality estimation. A) Graph shows intrinsic dimensionality (as measured by correlation dimension) computed on EMT data for different amounts of diffusion time (t) for three values of adaptive kernel *(ka =* 4, 10, 30). The peak values suggest optimal diffusion times that restore maximal dimensionality (information) to the data. B) 3D scatter plots of E-cadherin, Vimentin and Fibronectin, colored by Zeb1 for different values of *t* and *ka*. Scatter plots from the optimal *t* for each *ka* are outlined with green boxes. We see that the relationships have most detail and are most informative at the optimal times. C) Intrinsic dimensionality versus diffusion time graph for bone marrow data (from Fig. 3) for adaptive kernel values (*k_a_* = 4, 6, 30). D) 3-D scatterplot of Cd34, Cct5 and Pf4 colored by clusters from (Paul et al. 2015), the relationships at optimal diffusion times are outlined in green. The relationship and trajectory structure shows most detail at the optimal diffusion times

### Data preprocessing

MAGIC can be generally applied to any type of high dimensional single cell data to remove noise and clarify structure in the data. However, before a cell-cell distance matrix is computed, each data-type typically requires data specific pre-processing and normalization steps. Pre-processing is particularly important in the case of scRNA-seq, to ensure that distances between cells reflect biology rather than experimental artifact. We perform two operations on the data which are typically applied to this data type: 1) library size normalization on the cells, and 2) Principal Component Analysis (PCA) on the genes.

scRNA-seq data entails substantial cell-to-cell variation in *library size* (number of observed molecules) which is largely due to technical variation occurring due to multiple enzymatic steps, such as lysis efficiency, mRNA capture efficiency and the efficiency of multiple amplification rounds(*15*). For example, the cell barcode associated with each cell can have a substantial effect on the PCR efficiency and subsequently the number of transcripts in that cell.

Thus, some cells are highly sampled with many transcripts, while other cells contain far fewer transcripts. However, this variation is often technical and not reflective of underlying biological differences in cell size. Therefore, we normalize transcript abundances (library size), so that each cell will have an equal transcript count.

Given a *m * n* data matrix *D,* the normalized data matrix is defined as follows:

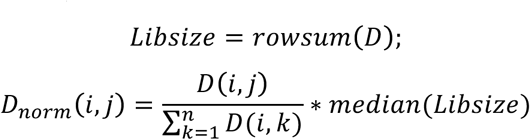

This effectively eliminates cell size as a signal in the measurement for the purposes of constructing the affinity matrix and thus the resulting weighted neighborhood is not biased by cell size.

Second, we perform Principal Component Analysis to further increase the robustness and reliability of the constructed affinity matrix. While drop-out renders single cell RNA-seq data extremely noisy, the modularity of gene expression provides redundancy in the gene dimension which can be exploited. Therefore, we perform PCA based dimensionality reduction to retain ~70% of the variation in the data, which typically results in 20 to 100 robust dimensions for each cell. Then, the cell-cell affinity matrix is computed off of these PCA dimensions.

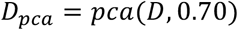

However, once *M^t^* is computed, we have a vector of weighted neighbors associated with each cell in our data. We can now use this robust neighborhood operator to impute and correct data using the library-size normalized count matrix (before PCA). Thus, while we use PCA to gain more robustness for the computation of *M,* the imputation *D_imputed_ =M^t^ * D* is performed at the resolution of individual genes.

### Validating MAGIC using synthetic “ground truth” datasets

We evaluated MAGIC on 3 different scRNA-seq datasets comprising different biological systems and measurement technologies and in each case demonstrated that MAGIC successfully recovers known biological relationships that were indiscernible prior to imputation. This provides a qualitative assessment of MAGIC, but fails to provide a quantitative quality metric. Such quantification requires ground truth for direct comparison with the imputed data. However, due to technological limitations, there is no single cell transcriptomic dataset that does not suffer from the very dropout MAGIC was designed to correct. Instead, to quantitatively evaluate the accuracy of MAGIC's imputation, we construct two semi-realistic datasets, where a ground truth is known prior to a synthetic dropout procedure.

First we created a validation dataset was based on bulk transcriptomic data from 206 developmentally synchronized C. elegans young adults, measured using microarrays (*35).* These samples are taken at regular time intervals during a 12-hour developmental time-course and consecutive samples exhibit relatively small changes in expression. While these samples do not come from single cells, as a developmental trajectory, this data is high dimensional in measurement space, but the data lie on a lower dimensional manifold. Since these are bulk measurements, we expect most related genes to be detected in each sample.

We down-sampled this data to emulate the sparsity found in scRNA-seq data. Since the data was log-scaled, we first exponentiated the data. Then we down-sampled each gene by subtracting random values sampled from an exponential distribution, such that 0%, 60%, 80% and 90% of the values are 0 after down-sampling (see Figure 8A). Then we log-scaled the data again and normalized each gene using a z-score. To evaluate the accuracy of MAGIC's imputation we applied MAGIC (with parameters *npca=20,* ka=3, *t=5)* to this synthetically "dropped out” data and then compared between the original and imputed data. We note that this dataset is particularly challenging as it only contains 206 samples, whereas MAGIC is primarily intended for datasets consisting of thousands of samples, as is the case for most single cell datasets. With few data points, MAGIC can recapitulate major trends that span across the data. Therefore, similar to the analysis performed in the original publication of the data, we select only genes that load to the first two PCA components of the data. This results in a data matrix with 206 worms and 9861 genes.

**Figure 8:**
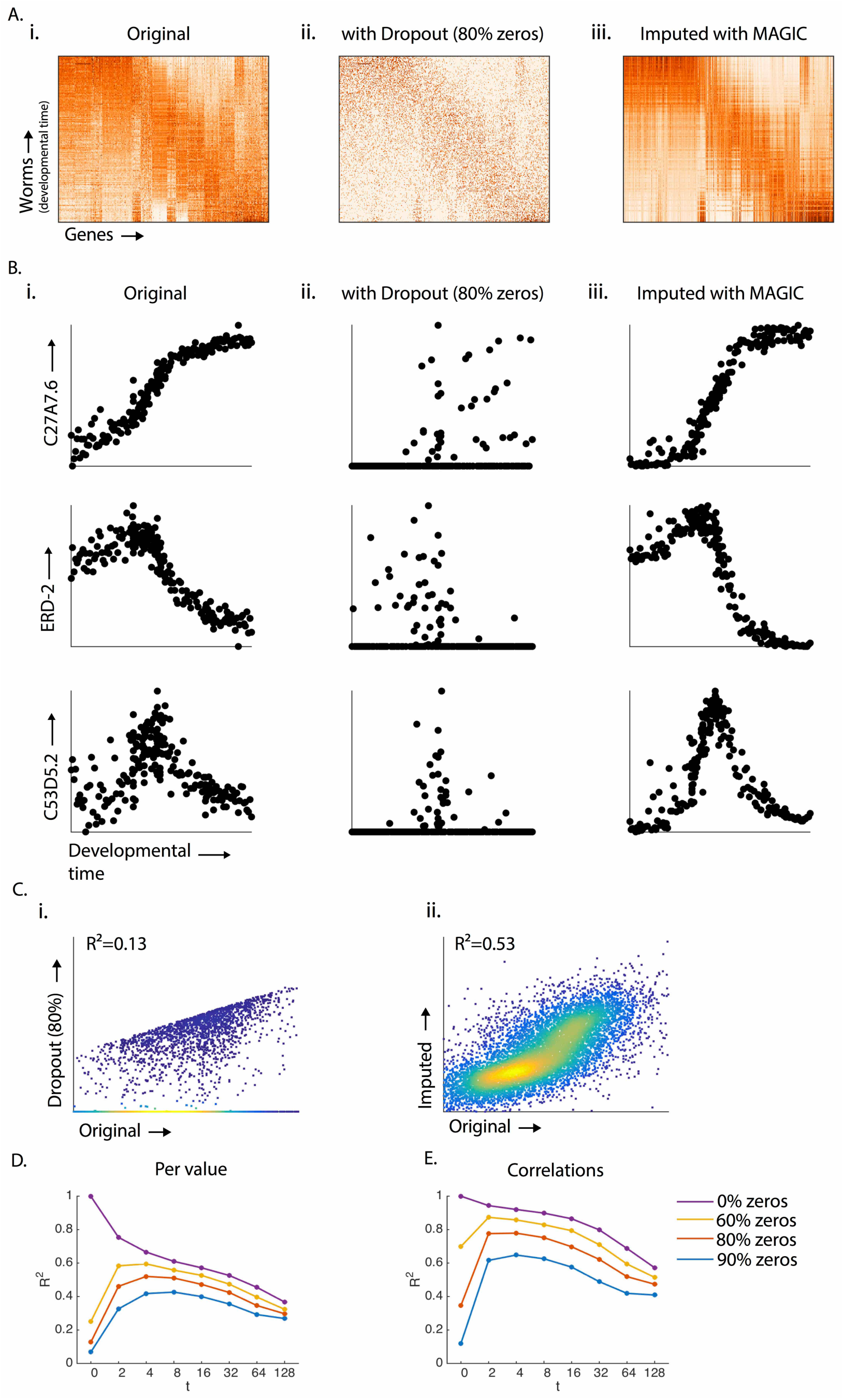
Validation of MAGIC using bulk C. elegans gene expression data with artificially induced dropout. A) The gene expression matrix with 206 worms sorted by developmental time along the Y-axis, and genes (along columns) clustered hierarchically. (Ai) The original matrix, (Aii) the matrix after 80% of the values are set to 0, and (Aiii) restored values after MAGIC. B) Scatter plots of gene expression (Y-axis) as a function of developmental time (X-axis) for three genes: C27a7.6, Erd-2 and C53d5.2. Bi) the original gene expression vs time, Bii) gene expression after dropout, Biii) after magic (with diffusion time *t*=5). Ci) A scatterplot of values comparing the data before (y-axis) and after (x-axis) dropout. R^2^=0.13 quantifies the discrepancy between values after dropout. Cii) A scatterplot of the original values (y-axis) and values after MAGIC (x-axis) with diffusion time *t*=5. R^2^=0.53 quantifies the recovery of values after imputation. D) Line plots showing the recovery of values (R^2^ of imputed values with original values) after MAGIC at various diffusion times. The different curves show recovery for different levels of dropout (purple=0%, yellow=60%, red=80%, blue=90%). E) Shows line plots quantifying the recovery of gene-gene correlations after MAGIC with various diffusion times. The original correlation matrix is compared to the imputed correlation matrix and the match is quantified by R^2^.

We first qualitatively assess MAGIC's ability to recover key patterns and trends in the data. Looking at the expression matrix, the imputed data largely matches the same structure and trends observed in the original data (Figure 8A). In fact, the pattern presented in the imputed data seems cleaner and sharper than that in the original data. We note that the original data was collected in 2002 (*36*) with early (noisy) microarray technology. Hence, it is quite possible that the data after imputation might be a less noisy version of the true underlying biology. To zoom in onto finer structure and illustrate MAGIC's ability to recover key trends in the data, we focus 3 selected genes (C27a7.6, Erd2 and Cd3d5.2) based on their non-monotonic developmental time trends and compare the original and imputed shapes for each of these trends. Indeed, for each gene we find close concordance in the developmental trend between the original and imputed data (8B). We note that while ~80% of the values are zero in the dropped-out data, all these follow a smooth developmental curve following imputation. We see a monotonic increase in C27a7.6 (a membrane transport protein), decrease in Erd2 (a homolog to the human ER protein retaining receptor), and non-monotonic curve in Cd3d5.2 (a gene predicted to have zinc ion binding activity). Overall, MAGIC performs very well in recapitulating the ground truth even under a severe degradation of the dataset.

The real advantage of having ground truth is the opportunity to quantitatively evaluate MAGIC's accuracy by directly comparing the original and imputed values. Figure 8C shows the scatter plot comparing each entry of the original matrix and the matrix with 80% zeroes. The overall correlation of values (each entry of the data matrix) before and after dropout is only R^2^=0.13. However, after MAGIC, we see this value substantially increased to *R^2^=* 0.53. An important point to note regarding the imputed data is that while MAGIC does not fully restore the original data, it rarely imputes values that do not exist in the original data matrix. Figure 8D shows a more comprehensive evaluation of the correlation between the original and imputed values for various values of dropout and values of *t* (time of diffusion). For example, at dropout of 90%, the R^2^ increases from 7% to 43% (at *t*=8).

One of our main goals in developing MAGIC was to be able to capture gene-gene relations from single cell data. Hence it is important to show that MAGIC not only captures univariate features of cells, but can also capture multivariate relations effectively. We evaluate MAGIC's ability to capture gene-gene correlations by comparing these correlations between the original and imputed data. Surprisingly, the agreement between the original and imputed data is even higher in the case of gene-gene correlations, than that of the univariate case (Figure 8E). For example, the agreement in gene-gene correlations between the original data and data with 90% of the values dropped out is 0.12. MAGIC recovers most of the gene-gene correlations so that after imputation we have a *R^2^* of 0.65. For 80% of the values at zero, MAGIC improves from 0.35 to 0.78. This is likely because genes that have good correlations between them are more structured in the data and MAGIC can then use this structure for its recovery. Therefore, we conclude that MAGIC can be an effective tool for exploring gene-gene relations.

We found evaluation of the worm data particularly encouraging, considering both the noise levels typical of microarray data and that only 206 data points were included, far less data points than in typical single cell datasets. We therefore wanted to also evaluate a dataset with more data points. We used the MAGIC-imputed count matrix of the EMT data presented in Figure 4 as the "ground truth” of a synthetically created dataset and then re-created synthetic dropout (0%, 60% 80% or 90% zeros) of this "imputed ground truth”. We re-impute the data using MAGIC (setting *ka*=10 and *t*=[0, 2, 4, 6, 8, 16, 32, 64, 128]).

Indeed when MAGIC is supplied with >7000 cells, imputation becomes substantially more accurate (Figure 9A). With 90% zeros, the R^2^ with the original data is brought down as low as the worm data, 4%. However, after MAGIC the R^2^ between the imputed and original data goes up to 70%. Similarly, we see that with 80% zeros, we have 9% after dropout, which is corrected to 81% after imputation. For 60% zeros, we have 21% after dropout, which is corrected to 88% after imputation. While these values are very encouraging, we note that this synthetic data is likely less noisy and therefore easier than a typical single cell dataset. Nevertheless, they demonstrate very powerful imputation and recovery for a test dataset.

**Figure 9:**
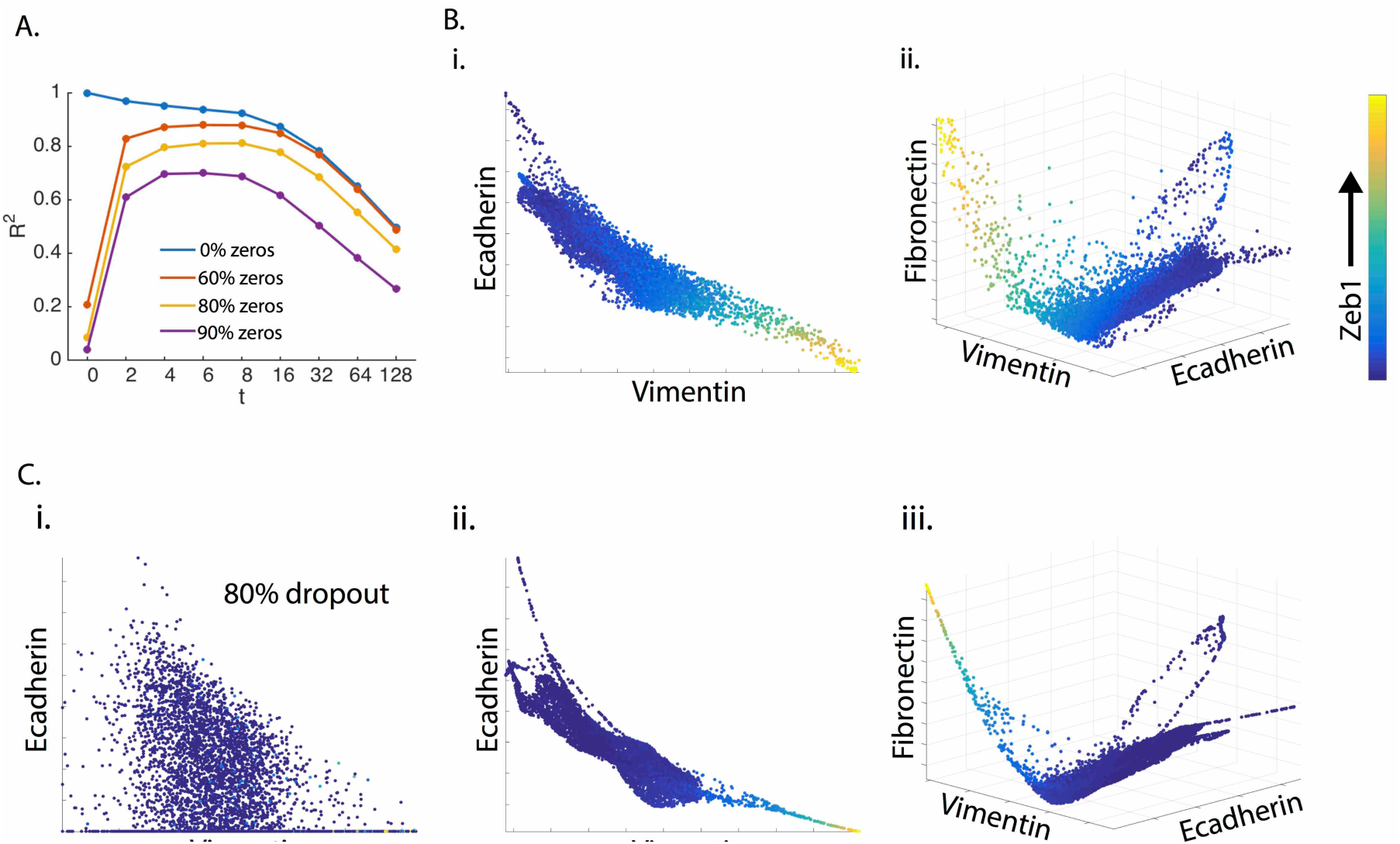
Validation of MAGIC using synthetic EMT dataset. A) Line plot shows recovery of values quantified by *R*^2^, for different levels of artificially re-induced dropout (blue=0%, red=60%, yeNow=80%, purple=90% zeros) for various diffusion times. Bi) 2D scatter plot of canonical EMT genes E-cadherin and Vimentin, colored by Zeb1 Bii) and 3D scatter plot of Ecadherin, Vimentin, Fibronectin, colored by Zeb1 before dropout. C) Shows the dropped-out and recovered versions of the plots in (B). Ci) Ecadherin and Vimentin after 80% of values set to 0. Cii) Scatterplot of recovered Ecadherin and Vimentin. Ciii) 3-D scatterplot of Ecadherin, Vimentin and Fibronectin after 80% dropout and recovery.

An important feature of MAGIC is that it is particularly good at capturing the "shape” of the data. Comparing the key edges in the original data (9B) with the imputed data (9C) we observe recovery of all key structural features of the data. We do note that the imputed data is less noisy and more accurately adheres to a low dimensional manifold. We conclude that in addition to removing technical noise, MAGIC likely removes true biological variation, and the resulting imputed data likely under-estimates the natural "biological noise” in the system.

### Robustness of MAGIC to Subsampling

An important feature of any algorithm is its robustness to input parameters and subsampling of the data (in this case, cells). Figures 8 and 9 already demonstrated the importance of having a robust number of cells for effective recovery of the data.

First, we consider sensitivity of MAGIC to subsampling of cells. We start with the 7523 cells collected in the EMT HMLE data of Figure 4 and consider the imputation result on the full data as the ground truth. For this analysis, we only consider the 9,571 genes that are expressed in more than 250 cells, to ensure these genes will likely remain present in each subsample. More generally, we expect the quality of the imputation to depend on gene expression, both the absolute expression level of a gene when it is observed, as well as how frequently (in how many cells) it is observed. To take this into account, we divide remaining genes into two groups, based on the mean log expression in the raw data, highly expressed genes (3,190 genes) and lowly expressed genes (6,381 genes). We subsampled cells to different degrees, uniformly at random (100 iterations each). For each subsampled dataset, we remove any genes that have no expression and impute the remaining genes using MAGIC (for the same set of parameters). For each imputed matrix, we compute the correlation-squared, R^2^, per entry against the ground truth (full dataset). Figure 10Ai shows the mean correlation-squared across 100 iterations with 1-standard deviation represented by the error bars. MAGIC is highly robust to subsampling of cells across both groups of genes. Even for a subsample with only 1000 cells, we obtain R^2^ > 0.94 among highly expressed genes and R^2^ > 0.61 among lowly expressed genes (with standard deviation < 0.01 for both).

**Figure 10:**
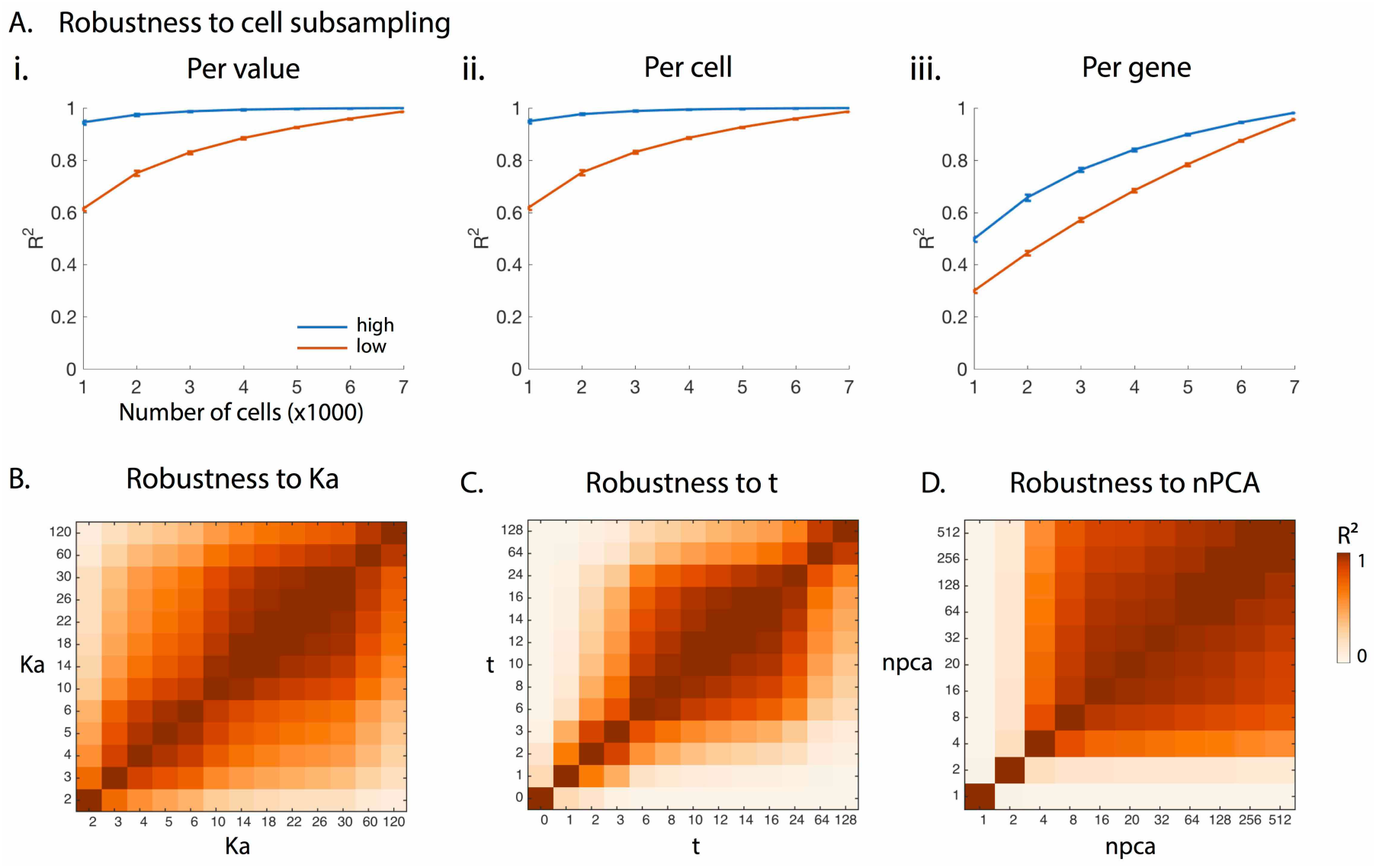
Robustness of MAGIC to cell subsampling and parameters. A) *R^2^* of original to re-imputed values on 9,571 genes that are expressed in more than 250 cells and divided into two groups based on expression levels (blue = 6381 high expressing genes, red = 3190 low expression). The *R^2^* of (i) values, (ii) cells, (iii) genes (original vs imputed) is computed for different levels of cell subsampling, with 100 random trials for each level of subsampling. The line plots show average and standard deviation of the *R*^2^ between full and subsampled data. B) The robustness of MAGIC to the adaptive kernel being set to various values for the *kath* nearest neighbor. The matrix shows the *R^2^* between various settings of *ka* with *t=6* and *npca*=20, indicating large ranges of high recovery stability. C-D) Similarly show robustness to diffusion time *t* and number of pca dimensions used in data preprocessing before distance and affinity computations respectively

Since our main interest lies in the quality of imputed cells, for each imputed cell (represented as a vector of gene counts) we compute the correlation-squared, R^2^, against the ground truth for the same cell and average the result over all cells. This "cellcentric” view of the data (Figure 10Aii) produces the same results and quality as the correlation observed across the full matrix. As demonstrated in previous analysis, MAGIC learns a lower dimensional manifold where cells reside and inferred cells adhere to this learned structure.

However, a "gene centric” view of each imputed gene (represented as a vector of cells), gives slightly different results (Figure 10Aiii). While we have good agreement when large numbers of cells are subsampled, e.g. when sampling 5000 cells averaged over all genes, R^2^ > 0.89 (std. < 0.01) on the set of highly expressed genes and R^2^ > 0.78 (std. <01) for the lowly expressed genes. This correspondence declines linearly with the number of cells subsampled, so that with only 1000 cells, we find R^2^ > 0.49 (std. < 0.01) on the set of highly expressed genes and > 0.29 (std. < 0.01) for the lowly expressed genes. Most genes are only observed in a fraction of cells, thus as the number of cells decline, so does the number of observations we have for any given gene. We find that we are successful at inferring genes that have high loadings on to the top PCA (or diffusion) components. That is, some genes behave in a more structured manner, and MAGIC is good at inferring these genes. But, not all genes exhibit such structured expression. Importantly, we have the ability to predict in advance (based on their PCA loadings), which genes we are likely able to impute well.

### Robustness of MAGIC to Parameters

MAGIC requires three key input parameters, *ka* (to set the adaptive kernel to the distance of the kth nearest neighbor), *t* (the number of times M is powered) and *npca* (the number of PCA components used to construct the affinity matrix). While we proposed criteria to guide the choice of these parameters, we also analyze MAGIC's robustness to their exact values.

MAGIC uses an adaptive kernel for cell-cell affinity computation, where σ, the width of the Gaussian kernel at each point is set to the distance to its k^th^ nearest neighbor (denoted *ka* (“adaptive k”)). We generally pick *ka* such that it is the smallest value that still results in a connected graph. We test MAGIC's robustness to *ka,* applying a range of *ka* values to the EMT data presented in Figure 4, with *t* set to 6 and *npca* to 20. To avoid the possibility of correlation being dominated by a small number highly expressed genes, we use z-score values for each gene in the imputed matrix. Then, we compute the R^2^ of the post-imputation data for each pair of *ka* settings (Fig 10B). MAGIC is highly robust for a suitable range of *ka* values (between 10-30), the average R^2^ value for *ka* = 10-30 is 0.95 (std 0.05). However, a very large value of *ka* (60-120) *over-smooths* the graph resulting in a weaker correlation score with other settings of *ka* (mean 0.56, std 0.27).

Next, we consider robustness of MAGIC to the diffusion time (t), by applying MAGIC to a range of values, keeping other variables fixed *(npca=20, ka*=10, Figure 10C). Again, we find that MAGIC is robust to a suitable range of *t* (6–24). In particular, the average R^2^ value for *t*=6-24 is 0.90 with a standard deviation of 0.10. However, a very large value of *t* (64-128) *over-smooths* the graph resulting in a weaker correlation.

Lastly, we consider robustness of MAGIC to the number of PCA *(npca)* components used to build the affinity matrix. We compute MAGIC based on a range of values of *npca,* holding other parameters fixed *(k_a_*=10, *t*=6). As shown in Figure 10D, we find that MAGIC is highly robust to the choice of *npca.* In particular, for *npca>=* 16, the average R^2^ is 0.94 with a std of 0.05. However, as expected, since few number of PCA components do not capture enough variance in the data, we observe low correlation between small and high *npca.* Overall we conclude that MAGIC is robust to a wide range of parameters, around the level that our heuristics for *ka, t* and *npca* provide. Thus changes in these parameters should have minimal effect on imputed results.

### Gene-Gene Interactions with KNN-DREMI

Our motivation for developing MAGIC was to recover gene-gene relations from sparse and noisy single-cell RNA-seq data. We have demonstrated that while relations are often obscured due to dropout (Figure 1B), MAGIC enables detection of gene-gene interactions system wide (Figure 1C, 2E-G, 4D, 5C). Thus, after MAGIC is applied to the data, gene-gene interactions can be clearly investigated using 2D or 3D biaxial plots, for an unbiased genome-wide exploration. However, it is impractical to plot and eyeball biaxial plots for all pairs of genes (nearly 1 billion pairs for a genome-wide measurement). We would thus like to automatically score gene-gene relationships to find regulators of genes and targets of regulators, and to infer large-scale regulatory networks. Moreover, the observed relationships are often highly non-linear and thus we need a shape agnostic metric, even where simple correlations fail to capture the trend. Therefore, we adapt our previously published DREMI method (conditional-Density Resampled Estimate of Mutual Information), a shape-agnostic information theoretic measure to compute the strength of gene-gene interactions (*11),* to scRNA-seq data. DREMI was well suited for mass cytometry data, which involved a lower dimensionality (30-50 dimensions) and a substantially larger number of observed cells. We modify DREMI to adapt effectively to the higher dimensionality and cell sparsity of single-cell RNA sequencing data. The key change is the replacement of the heat diffusion based kernel-density estimator from (*37*) by a k-nearest neighbor based density estimator(*38*). We term this metric *k*NN-DREMI.

The main idea underlying the DREMI method is the use of conditional density instead of joint density in computing the "information” that is shared between two variables. In single-cell data we found that most cells resided in one or two dense areas, where as most phenotypes (and often the most interesting ones) were only sparsely populated. Mutual information, which weights based on density, is largely dominated by small dense regions and hence does not capture the relationship across the full dynamic range of the data. Moreover, differences between cells within these dense regions are often driven by measurement noise, rather than then real biology, rendering mutual information an ineffective metric to quantify the relationship between two molecules. By using conditional density, DREMI better captures the functional relationship between two molecules across their dynamic range. DREMI first estimates a kernel density estimate (KDE) to smooth and fill in gaps in the data(*37, 39, 40*). Kernel density estimation involves placing a smooth kernel (such as a Gaussian kernel) at each data point and summing these Gaussians along a regular grid. Then DREMI normalizes the estimate along one of the two dimensions to form a conditional density estimate. The main steps of DREMI include:

1. Kernel density estimation to compute *p(x,y)* for two variables x and *y*.
2. Coarse-graining of KDE into larger discrete bins for entropy computation.
3. Normalization of the coarse-grained KDE to compute 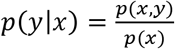.
4. Entropy and mutual information computation based on the discrete bins.

The main problem with applying this approach to single-cell RNA-seq data is that the typical computation of kernel density estimation does not scale well to higher dimensions and sparse data. In *k*NN-DREMI we use a *k*NN-based density estimator (*38*), which has been shown to be an effective method for sparse and high dimensional datasets. This involves a local computation involving only the k-nearest neighbors for each cell, which scales linearly with the number of cells. Moreover, while density estimation becomes prohibitively slow at higher dimensions and requires exponentially more data for stable estimates (*41*), a neighbor-graph has no dimensions and is only dependent on the computation of a good affinity matrix. The steps of *k*NN-DREMI are explained below:

#### 1) Computation of joint density using a *k*NN-method

Figure 11A shows the computation of a 2-dimensional joint density estimate using a *kNN* approach. The joint density is computed on a fine grid of points (shown in gray). Each grid point obtains its density from its kth nearest data-point neighbor. Figure 11Ai shows two data points colored by density based on their distance to the nearest neighboring datapoint (*k*=1). Thus the density at each grid point is calculated by by:

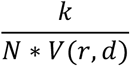

**Figure 11.**
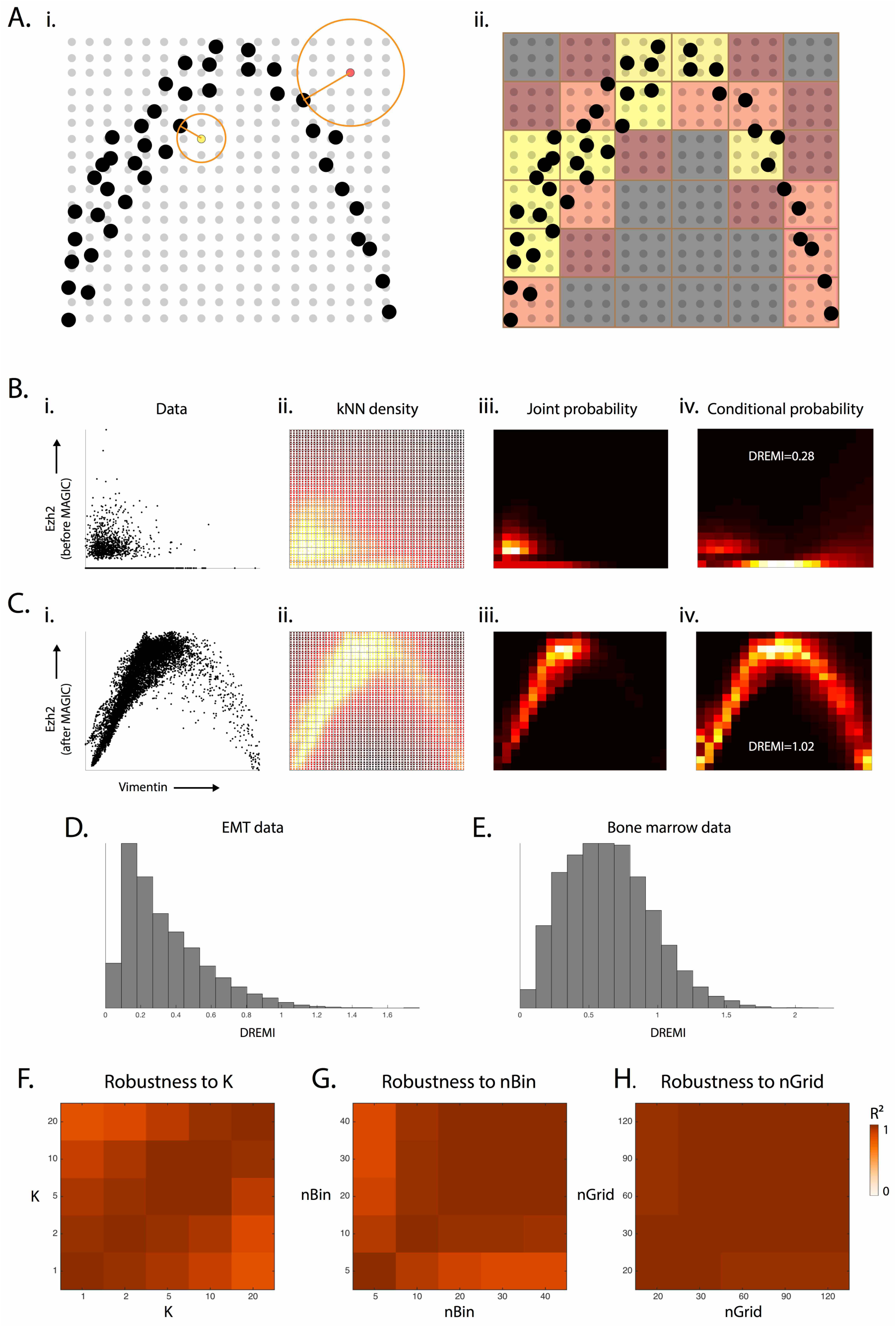
Computation steps of *k*NN-DREMI. Ai) Illustrates the computation of kNN-based density estimation on an 18×18 grid, shown as gray points with data points shown in black. Density is estimated at each grid point (yellow, and red grid points are examples) as inversely proportional to the volume of a circle with radius R equal to the distance to its nearest data neighbor (black point). Aii) After density estimation on the grid-points, the grid is coarse grained into a 6x6 discrete density estimate (red and yellow squares show coarse grained partitions) by accumulation of all densities within each square bin. B) The steps for computing kNN-DREMI are shown for Ezh2 (Y-axis) and Vimentin (X-axis) before MAGIC (ii) Scatter plot of the data, (ii) kNN-based density estimation on a fine grid (60×60) (iii) coarse-grained joint probability estimate on 20x20 partition (iv) normalization of joint probability to obtain conditional probability density, resulting in *k*NN-DREMI = 0.28. C) Same steps as (B) shown after MAGIC resulting in a *k*NN-DREMI= 1.02. D) Histogram of *k*NN-DREMI values on EMT datas. E) Histogram of *k*NN-DREMI values on bone marrow data. F-H) Show robustness of *k*NN-DREMI estimates to a range of values for *k*, *nBin* and *nGrid* respectively. F) Heatmap of correlation matrix where each entry shows the correlation between *k*NN-DREMI values on 3000 pairs of randomly chosen genes, for a pair settings for k, the nearest neighbor cardinality for the density estimate of each point. G) Heatmap of correlation matrix for 3000 randomly chosen edges for each pair of settings for *nBin,* number of bins used for coarse graining density estimate and computation of mutual information, H) Similar matrix as F,G for *nGrid,* the grid size on which *k*NN-based density estimate is computed for *k*NN-DREMI.

Where the volume of a d-dimensional ball of radius r is given by:

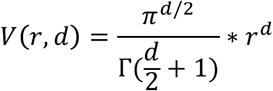

Since we are computing pairwise relationships, d=2 in this context. We set k=10 for robustness against outliers.

#### 2) Coarse graining the density estimate

While the KDE is computed on a fine-grid, the discrete mutual information is computed using few larger partitions (Shown in Figure 11Aii as square partitions). There are two reasons for this: first the scale of the density estimate is not necessarily the appropriate scale to compute mutual information. The density estimate is intended to smooth and fill in gaps in the data distribution and therefore requires a finer scale of resolution. The mutual information can be computed at a coarser scale to find clear and strong relationships. Second, having a coarser-scale resolution for mutual information renders the mutual information more robust, and less dependent on noise in partitions and lack of samples due to sampling irregularities. Therefore, we use on the order of 20 partitions in both the X and the Y directions and accumulate the density estimates for each grid point in the partition to obtain the coarser joint density distribution on which to compute entropy (Figure 11Aii).

#### 3) Computation of conditional density using a *k*NN-method

The key insight in DREMI is the increased fidelity of computing relationship strength on the conditional density rather than joint density to account for differences in sampling. For instance, in the relationship shown in Figure 11c,i, we see that the left half of the relationship is much more densely sampled than the right half and that the joint density (shown in Figure 11c,iii) only picks up signal in the left half. The mutual information, based the joint density, will only score the left half of this relationship. By contrast, the conditional density estimate (shown in Figure 11c,iv) picks up the relationship in both halves revealing a quadratic relationship.

To compute the conditional density estimate, we simply column-normalize the joint density estimate, i.e., divide the joint density estimate by the marginal. More formally, in the two variable case, if the joint density estimate is performed on a *η * n* matrix G, to condition on the second dimension, divide each entry by the column-total:

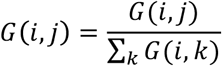

#### 4) Computation of Mutual Information from conditional density

The final step of *k*NN-DREMI is the computation of entropy and mutual information using the coarsegrained conditional density estimate from step 3. In the discrete case where *X* and *Y* can take on values between *1* and *m*, mutual information between two variables *X* and *Y* is generally computed as the difference between the entropy of *Y*, and its conditional entropy after conditioning on *X*:

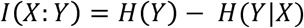

Here *H* is the Shannon Entropy is:

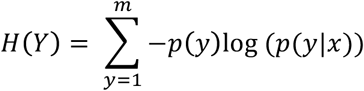

Conditional Shannon Entropy is given by:

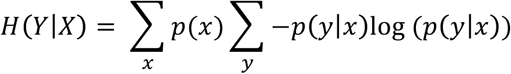

After computation of the coarse-grained conditional density estimates, we simply compute the mutual information using the equation above. Effectively, this simply added another level of conditioning to the original formulation of mutual information:

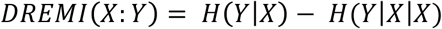

We illustrate this computation in Figure 11B-C on the relationship between Vimentin and Ezh2. Figure 11B shows the same data before MAGIC and Figure 11C shows this data after MAGIC, revealing clear non-linear relationship between the two variables, with Ezh2 peaking at intermediate levels of Vimentin.

Figures 11B,Cii, show the *k*NN-based joint density estimation performed on a fine grid (60 × 60). Each grid point gets density inversely proportional to its nearest data neighbor. Figures 11B,Ciii show the coarse-grained joint density estimate on 20 larger partitions. This joint density estimate is then converted into a conditional density estimate by column normalization in Figures 11B,Civ. This shows the boost in the signal in the second half of the relationship. Finally the *k*NN-DREMI score is computed as mutual information on this conditional density estimate.

The relationship between Vimentin and Ezh2 (Figures 11B,C) gets a *k*NN-DREMI score of 0.28 and 1.02 before and after MAGIC respectively. To place these scores into perspective Figures 11D,E show histograms of *k*NN-DREMI scores of 10,000 random gene-gene relationships in the EMT (Figure 11D) and bone marrow data (Figure 11E). We can see that 0.28 is not a high score while 1.02 is high on this scale.

*k*NN-DREMI requires three parameters to compute the DREMI score between two gene expression vectors; the number of neighbors for *kNN* density estimation (*k*), the size of the fine-grained grid on which *k*NN density is computed *(nGrid—square root of square grid size),* and the number of bins per dimension for course graining the densities *(nBin).* We choose *k* such that it computes density locally, thus a small number. We set it to 10 to increase its robustness, e.g. *k*=1 could be an outlier. *nBin* should be chosen such that enough resolution exists to accurately compute mutual information, but small enough such that each bin has a fairly large amount of data points. We set *nBin* to 20, thus giving 400 2D bins. Finally, *nGrid* should be significantly larger than *nBin* such that multiple grid points exist within each course bin. As a rule of thumb we set this value to 3 times *nBin,* thus 60. While our parameter choices are based on reason, there may be concern that the parameters have a large effect on the DREMI score. To investigate this we performed a robustness analysis to changes in the three parameters (see Figure 11F-H). We computed *k*NN-DREMI for 3000 random gene pairs of the EMT data for the following parameter values: *k* = [1 2 5 10 20], *nBin* = [5 10 20 30 40], and *nGrid* = [20 30 60 90 120], around the default parameter setting *k*=10, *nBin=20* and *nGrid=60.* To quantify robustness we computed R^2^ between each pair of parameter settings, for each of the three parameters. Figures 11F-H show that the *k*NN-DREMI score is highly robust to changes of the parameters within a reasonable range.

### Gene-Gene Interactions in EMT and Bone Marrow Datasets

Figure 12 shows gene-gene interactions from the EMT dataset (Figures 12A-C) and Bone marrow dataset (Figure 12D) after MAGIC. Here we see that MAGIC is able to impute clear, and often non-linear trends, even from from sparse data. Further, we see that these shapes can be scored by *k*NN-DREMI successfully. Figure 12A shows that MAGIC substantially increases our ability to detect gene-gene interactions, while the pre-MAGIC DREMI range is between 0-0.4, after MAGIC, this range increases to 0-1.7, with the mode shifting from 0 to 0.2. We note that there is almost no correlation between the DREMI scores before and after MAGIC and moreover, we find gene pairs with very high-DREMI after MAGIC, across the entire range of DREMI scores before MAGIC.

**Figure 12.**
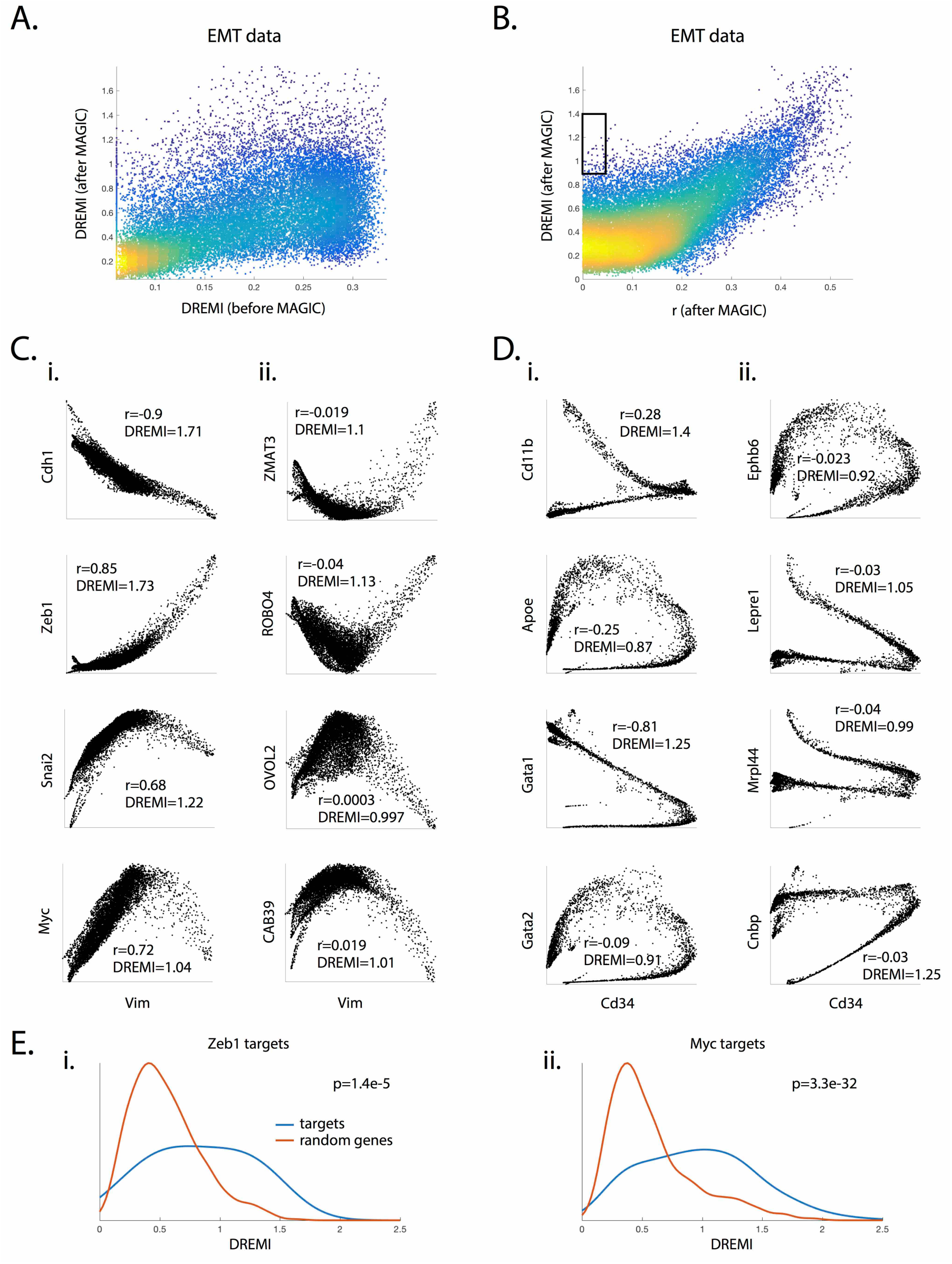
*k*NN-DREMI uncovers new and non-linear gene-gene interactions. A) kNN-DREMI before (X-axis) and after (Y-axis) MAGIC colored by density. Each point represents one of 28910 relationships between Vimentin (as a measure of EMT progression) and all other genes present in our EMT dataset. Scatterplot indicates a higher range of kNN-DREMI (0-1.6 vs 0-0.4) after imputation and values do not correlate well with their pre-MAGIC values. B) Pearson correlation computed on the same post-MAGIC gene-gene relationships versus kNN-DREMI. The black box indicates a region of low Pearson correlation but high *k*NN-DREMI (r<0.05, DREMI>0.9). C) 8 gene-gene relationships (4-well known, and 4 less known) from EMT data after imputation with Pearson correlation and kNN-DREMI scores, indicating that kNN-DREMI is much more effective at recognizing non-linear relationships. D) Similar analysis of bone marrow data (as presented in Figure 3). E) Shows the comparative kNN-DREMI distributions for the targets of transcription factors Zeb1 (Ei) and MYC (Eii), as shown in red against 1000 random non-targets, shown in Blue. In each case the p-value as computed by the ranksum test indicate significantly higher *k*NN-DREMI score for targets of the respective transcription factors.

Figure 12B shows a key advantage of using DREMI over Pearson correlation to quantify relationships. DREMI is able to correctly score the strengths of the highly non-linear relationships we frequently see between genes. We see that if the correlation coefficient is high then DREMI will also be high. However, there are additional relationships (highlighted in the box in Figure 12B) that only DREMI identifies. For instance, Figure 12Cii shows Vimentin versus Zmat3 (a p53 pathway gene known to be down regulated at the start of EMT). The correlation between the two markers is weak (R = 0.019), however the shape agnostic DREMI gives a higher score of 1.1 (see histogram in Figure 11D). In EMT-related edges (shown in Figure 12C) we see that Slug (Snai2), a well-known transcriptional regulator of EMT increases with Vimentin initially until it reaches a peak point, after which it is down-regulated. Therefore, this TF starts the transition but perhaps is not needed to maintain the mesenchymal state. Figures 12Ci and 12Cii show four well-known and less well known relationships that have high DREMI scores with Vimentin.

Figure 12Di,ii similarly show well-known and new relationships between Cd34, a progenitor marker, and various genes that mark developmental lineages in the bone marrow. For instance, Cd11b (Shown in Figure 12Di) increases with the myeloid lineage and decreases with the erythroid lineage. Note that correlation is particularly bad at these types of relationships, which diverge between cellular sub-populations. Correlation only scores this relationship 0.28 while DREMI scores it 1.41.

Finally, Figure 12E shows the ability of MAGIC combined with *k*NN-DREMI to infer gene-regulatory network structure without the need for perturbations. Here, we see that DREMI can predict targets for Zeb1 and Myc, two TFs involved in EMT. We computed DREMI scores between the these TFs and their gene target sets (RNAi for Zeb1 (*42),* Broad MSIGDB hallmark gene set for Myc) and compared this to DREMI scores between the regulators and 1000 random genes. Figure 12E shows these distributions with their corresponding Wilcoxon rank-sum test p-values. We find that DREMI is significantly higher for the targets of these regulators as compared to random non-targets.

### Comparison of MAGIC to Other Methods

We compare MAGIC to current state-of-the-art methods to fill in missing data and reduce noise, SVD-based low-rank data approximation (LRA)(*43*) and Nuclear-Norm-based Matrix Completion (NNMC)(*44*). Both methods have a low-rank assumption, i.e., like MAGIC, they assume that the intrinsic dimensionality of the data is much lower than the measurement space and utilize a singular value decomposition (SVD) of the data matrix. The singular-value decomposition of the data matrix, is a factorization of the form *D = UEV\** where *U* contains the left singular vectors of *D, V* contains the right singular vectors of *D,* and *E* contains the singular values along the diagonal.

The two methods we compare against MAGIC work as follows:

1. SVD-based low-rank data approximation (LRA)(*43*): This method for derives a low-rank approximation of a higher rank data matrix. After performing SVD, a lower rank version of *D, D_low_* is created by taking only the first *k* columns of *U* and *E* and only the first *k* rows of *V*\*. This is because the first singular vectors, like PCA vectors, explain a larger variation in the data, while the subsequent vectors may correspond to noise. Therefore, the elimination of the lower singular vectors effectively de-noises the data, albeit, only using linear directions of variation.
2. Nuclear-Norm-based Matrix Completion (NNMC)(*44*): This technique is designed to recover missing values in data matrices, which could potentially address the dropout issue. MNMC restores "missing values” so that the rank of the data matrix is not increased, as computed through a linear programming optimization. However, since minimizing the rank of a matrix is a non-convex optimization, they optimize a convex proxy for rank, which is the *nuclear norm (sum of all singular values)* of a matrix.

First, we compared the performance of the three techniques on a two-dimensional Swiss roll (See Figure 13). We added Gaussian noise along the Swiss roll (Figure 13A), and then embedded the Swiss roll into 5000 dimensions via a random QR rotation matrix.

**Figure 13.**
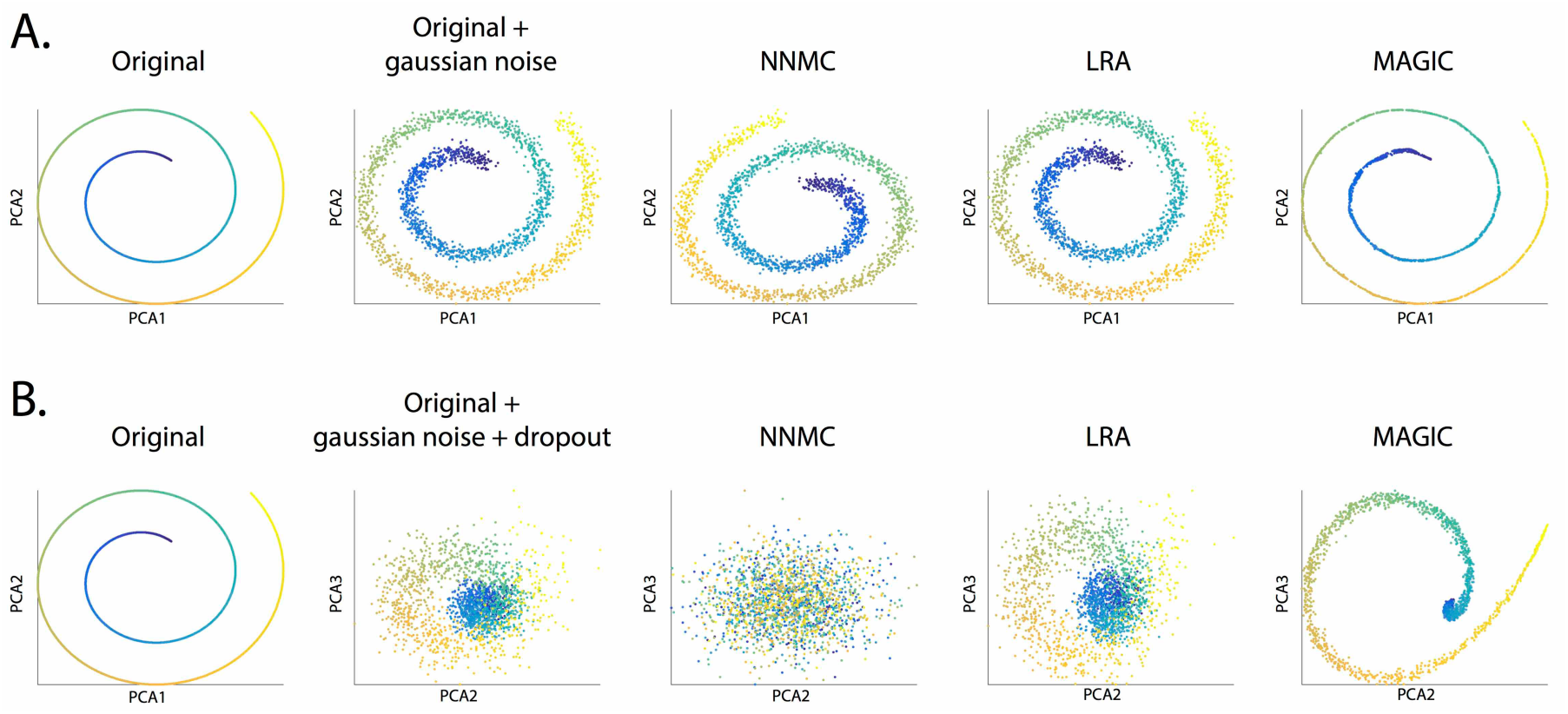
Comparison of MAGIC to other imputation methods on synthetic data. For all the plots in this figure, we use a 2-D Swiss roll, embedded in 5000 dimensions (by QR rotation) with Gaussian and dropout noise. A) First two PCA components of the noiseless (Ai) high-dimensional Swiss roll and Swiss roll after Gaussian noise are shown (Aii), (Aiii-v) Show same PCA components of the Swiss roll after imputation by nonnegative matrix factorization (Aiii), low-rank approximation (Aiv) and MAGIC (Av). Only MAGIC is able to reverse Gaussian noise along the Swiss Roll. B) Shows recovery after the addition of Gaussian noise along with random dropout. The dropout was also increased along the curve of the Swiss Roll. B) Recovery after Gaussian noise as well as dropout (Bii) is added to the Swiss Roll dataset after rotation into high dimensions. Biii-Bv) Show recovery after non-negative matrix completion (Biii), LRA (Biv) and MAGIC (Bv) respectively. Only MAGIC is able to recover the coiling shape of the Swiss Roll after dropout.

Results show that only MAGIC is able to denoise even relatively simple Gaussian noise. While LRA can take off noise only from outside the plane of the Swiss roll (by decreasing rank and essentially discarding noise dimensions), NNMC seems incapable of even that. NNMC is only concerned with retaining rank, and so it can fill in data arbitrarily so as not to increase rank.

The real advantage of MAGIC becomes clear when we add dropout, typical of scRNA-seq data (Figure 13B). Dropout was added to create 80% zeros, creating regions of different densities in the data. We find that only MAGIC is able to correct for drop-out and restore the Swiss Roll. The "recovered” LRA looks identical to the noisy, dropped out LRA, and the "recovered” NNMC looks cloud-like. We conclude that MAGIC is uniquely well suited to handle the drop-out rampant in scRNA-seq data.

We also compared all 3 techniques on 8 known biological relationships in our data (See Figure 14). In each case, NNMC performs poorly, generally only imputing a single linear shape. Occasionally the direction of correlation is also incorrect in NNMC. For, instance, the Cdh1 vs Cdh2 (E-cadherin vs N-cadherin) edge shown in Figure 14A, is known to have a negative relationship (*45).* However, NMMC imputes a positive correlation between these genes. Additionally, NMMC finds no relationship between the well-known negative correlation between canonical EMT markers E-caherin and Vimentin. A possible explanation for this poor recovery is that NMMC "trusts” non-zero values and only attempts to impute possibly missing zero values. Whereas in scRNA-seq drop-out of molecules impacts all genes and even non-zero genes are likely lower than their true count in the data. Hence NMMC is poorly suited to this data type.

**Figure 14.**
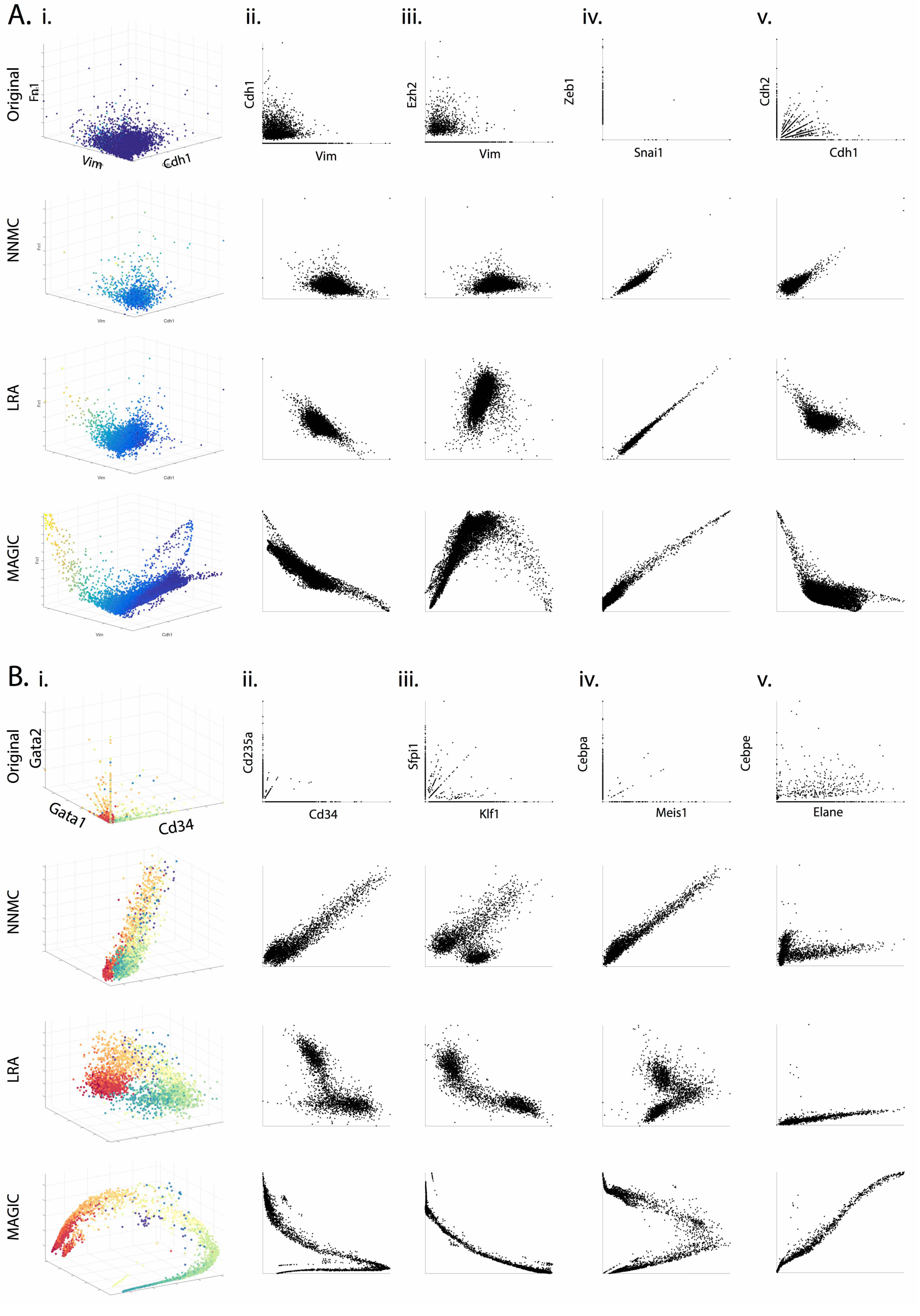
Comparison of MAGIC to other imputation methods on EMT and bone marrow data. A) Comparison of recovery with Non-Negative Matrix completion (NNMC)(*44*), Low-Rank Approximation (LRA)(*43*) and MAGIC on various edges of the EMT Dataset. The last row shows that the imputation by MAGIC gives the most detailed and non-linear interaction shapes. Ai) 3D Scatter plot of Vimentin, Ecadherin, Fibronectin, colored by Zeb1 after three methods of imputation. Only MAGIC is able to impute the finer branches leading to the stable mesenchymal state (high Zeb1) and apoptosis (low Zeb1). Aii) Scatterplot of Ecadherin vs Vimentin showing a non-linear negative relationship after MAGIC. Other methods are able to only catch slight negative correlation. Aiii) Ezh2 vs Vimentin: MAGIC is able to impute a negative quadratic relationship. Aiv) Linear relationship of Zeb vs Snail imputed by all methods. Av) Nonlinear negative relationship between N-cadherin and E-cadherin, mirrors the non-linear relationship between Ecadherin vs Vimentin as imputed by MAGIC. B) Comparison of imputation methods on gene-gene interaction recovery on the bone marrow dataset:.Bi) 3D scatterplot of Cd34 (a progenitor marker), Gata1 (a erythropoiesis marker) and Gata2 (a developmental marker) colored by cluster ID, shows MAGIC connecting clusters into an arc where only the erythroid branch shows increase in Gata1. Bii) Cd34 vs Cd235a (an erythroid marker) where MAGIC successfully imputes a negative relationship in only the erythroid branch of the data. NNMC imputes a positive relationship and LRA shows a fuzzy relationship. Biv) Cebpa (a myeloid marker). vs Meis1 (an erythroid marker) shows a negative relationship on erythrocytes as imputed by MAGIC and mild positive relationship on myeloid cells. Bv) Positive relationship between Cebpa and Elane (neutrophil markers).imputed by LRA and MAGIC, although MAGIC shows the more pronounced positive relationship expected for these markers.

LRA performs slightly better, as the most significant components of the SVD do usually contain the hyperplanes of the data manifold. However, it cannot separate the exact manifold from external noise, likely due to its inability to find non-linear directions in the data. Therefore, it cannot impute the fine-grained structure that MAGIC imputes as shown throughout Figure 14. For instance, in Figure 14Ai we see that MAGIC is the only method that is able to impute the details of the sparser branches, which contain the mesenchymal and apoptotic cells, while the other methods only impute a cloud shape. In the Bone marrow data shown in Figure 14B, we see that MAGIC is the only method that is able to clarify the developmental trajectory seen in 14Bi into an arc with myeloid cells developing to one arm and erythroid cells developing in the other arm.

## Discussion

Single-cell RNA sequencing is an emerging high-throughput technology that provides genome-scale transcriptomes of individual cells and therefore reflects their precise state, identity and potential activity. One of the most exciting potential uses of this data is system-wide mapping of regulatory relationships, based on naturally occurring variation between cells (*10-14*). Of great promise is the ability to infer such regulatory relationships in multiple primary cell types (e.g. whole tissue), including diseased tissues, without perturbation. However the drop-out rampant in this data limits our ability to interpret this data, and especially impedes our ability to identify the gene-gene relations.

Here, we presented the MAGIC algorithm to computationally overcome this challenge. MAGIC uses data diffusion to correct and impute missing single-cell RNA sequencing measurements. We provided extensive evaluation and validation of MAGIC and demonstrated that MAGIC is able to recover gene-gene interactions, developmental trends and enhance cluster-specific gene expression. Further, we presented an adaptation of the DREMI technique, termed KNN-DREMI, to quantify the strength of non-linear and noisy gene-gene relationships. MAGIC is able to impute back finegrained structure of gene-gene interactions including highly nuanced and non-linear shapes of interactions, whose strength can be successfully scored by KNN-DREMI. In addition to gene-gene interactions, these techniques can also be used to visualize relationships along temporal or pseudo-temporal trends in the data.

A number of recent publications (*46-48*) have demonstrated a powerful combination of CRISPR and single-cell RNA-sequencing that enable the reconstruction of genome-wide maps of regulatory relationships. However, these require extensive experimental work and perturbations, and as such, cannot be applied to primary tissue sources. MAGIC, enables the large-scale computation of pairwise gene-gene interactions from complex populations of primary cells without perturbation. This allows for the discovery of gene regulatory relationships in a wide diversity of primary cell types without perturbation. A possible application of this is to build an atlas of such regulatory networks in healthy individuals and compare these to their corresponding networks in diseased tissues. This can be applied to clinical material, such as biopsy samples, and help elucidate the regulatory relations that go awry in disease in a patient specific manner.

## Materials and methods

### Cell culturing and EMT induction

HMLE and all derived cell lines used in this work were cultured in MEGM (Mammary Epithelial Cell Growth Medium) media (Lonza, USA, CC-3051). Cells were cultured in round tissue culture dishes 10cm in diameter (Corning, USA) and split to a ratio of 1:7 every 2 to 3 days or once they reached 80% confluence on a plate. All cell dissociations were performed using TrypLE™ (Ambion, USA) reagent. EMT was induced by addition of Recombinant Human TGF-β1 (HEK293 cell derived) (PeproTech, USA 100-21) to a final concentration of 5ng/ml. All cells under induction were passaged once they reached 80% confluence.

### Indrop

scRNA-seq was performed using the Indrop protocol as described in (*49*). To prepare the cells for scRNA-seq experiments, they were cultured to 70% confluence and dissociated from the plate with the addition of 3ml of trypsin for 5 mins at 37 °C, After dissociation cells were kept at +4 °C at all times in MEGM-complete media. Two 1x PBS (Ambion, USA) washes were performed on the dissociated cells and cell viability was evaluated using trypan blue staining prior to scRNA-seq. All Indrop experiments were performed with cell viability exceeding 90%.

### Creation of the Synthetic Datasets

We created several synthetic datasets to demonstrate the effects of dropout, noise and recovery after application of MAGIC. These include:

1. Sigmoidal data: We generated a sigmoidal data 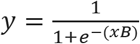 for 5000 points in the range −1 < *x <* 1 with *b* = 5. Then, we added Gaussian noise to the data, with mean 0 and standard deviation 0.1. This data is shown in Figure 1E and is colored by density. To simulate dropout, we randomly set 50% of the points to zero and for the remaining 50% of points we randomly removed *p%* of the values, where *p* was uniformly sampled between 0 and 100. The dropped out data is shown in Figure 1F.
2. Swiss roll data: To illustrate the MAGIC algorithm, we generated a Swiss roll dataset (Figure 2B). A Swiss roll is a prototypical example of a higher dimensional dataset with a continuous lower dimensional manifold. We first generated a 2-dimensional Swiss roll sampled at 1000 points. The data is embedded in 10 dimensions by random rotation via a randomly generated QR transformation. Then these 10 dimensions are extended to 100 dimensions by replicating each dimension 10 times with additional Gaussian noise. We added Gaussian noise with mean 0 and standard deviation 2.5. The first two PCA components of this data, illustrating the Swiss roll shape is shown in Figure 2B. For Figure 13A, the Swiss roll consisted of 2000 points. A Gaussian noise of mean 0 and standard deviation 0.35 was added to create a noisy Swiss roll. This was then embedded into 5000 dimensions via QR transformation. The first two PCA components of this data is shown in Figure 13A. In Figure 13B, we added dropout by subtracting values per data-point from an exponential distribution with *λ* = 10 in the inner part of the Swiss Roll and decreasing to *λ* = 1.5 towards the outer part of the spiral.
3. C. elegans bulk gene expression data with artificially-induced dropout: In order to validate MAGIC we utilized bulk microarray data of developing C. elegans embryos. Transcript abundances of 206 worms were used. In order to induce artificial dropout, the expression levels (which were log-scaled) were exponentiated, and then each entry was downsampled using an exponential distribution such that the result had 0%, 60%, 80% and 90% of the values set to 0. Then the data was log-scaled and normalized by zscoring. Additionally only genes that loaded significantly to the first two PCA components were utilized. This results in 206 worms with 9861 genes.
4. Recovered EMT data with artificially-induced dropout: We used the MAGIC-imputed count matrix of the EMT data as the "ground truth” of a synthetically created dataset and then re-created synthetic dropout. Starting with data from 7523 HMLE cells 8-10 days after TGFB treatment, we first imputed the data with MAGIC (*npca*=20, *ka*=10, *t*=6) and then we induce dropout by downsampling using an exponential distribution such that 0%, 60%, 80% and 90% of the values are set to 0.

## MAGIC Software

Python and matlab packages for MAGIC are available via github for academic use: https://github.com/pkathail/magic/.

## References

1. E. Z. Macosko et al., Highly Parallel Genome-wide Expression Profiling of Individual Cells Using Nanoliter Droplets. Cell 161, 1202–1214 (2015).

2. A. M. Klein et al., Droplet barcoding for single-cell transcriptomics applied to embryonic stem cells. Cell 161, 1187–1201 (2015).

3. K. Shekhar et al., Comprehensive Classification of Retinal Bipolar Neurons by Single-Cell Transcriptomics. Cell 166, 1308–1323 e1330 (2016).

4. A. Zeisel et al., Brain structure. Cell types in the mouse cortex and hippocampus revealed by single-cell RNA-seq. Science 347, 1138–1142 (2015).

5. P. Dalerba et al., Single-cell dissection of transcriptional heterogeneity in human colon tumors. Nature biotechnology 29, 1120–1127 (2011).

6. S. C. Bendall et al., Single-cell trajectory detection uncovers progression and regulatory coordination in human B cell development. Cell 157, 714–725 (2014).

7. L. Haghverdi, M. Buttner, F. A. Wolf, F. Buettner, F. J. Theis, Diffusion pseudotime robustly reconstructs lineage branching. Nat Methods 13, 845–848 (2016).

8. M. Setty et al., Wishbone identifies bifurcating developmental trajectories from single-cell data. Nat Biotechnol, (2016).

9. R. Satija, J. A. Farrell, D. Gennert, A. F. Schier, A. Regev, Spatial reconstruction of single-cell gene expression data. Nat Biotechnol 33, 495–502 (2015).

10. A. Ocone, L. Haghverdi, N. S. Mueller, F. J. Theis, Reconstructing gene regulatory dynamics from high-dimensional single-cell snapshot data. Bioinformatics 31, i89–96 (2015).

11. S. Krishnaswamy, Spitzer, M. H., Mingueneau, M., Bendall, S.C., Litvin, O., Stone, E., Pe'er, D., Nolan, G.P., Conditional Density-based Analysis of T cell Signaling in Single Cell Data. Science, (2014).

12. K. Sachs, O. Perez, D. Pe'er, D. A. Lauffenburger, G. P. Nolan, Causal protein-signaling networks derived from multiparameter single-cell data. Science 308, 523–529 (2005).

13. N. Rosenfeld, J. W. Young, U. Alon, P. S. Swain, M. B. Elowitz, Gene regulation at the single-cell level. Science 307, 1962–1965 (2005).

14. J. T. Gaublomme et al., Single-cell genomics unveils critical regulators of Th17 cell pathogenicity. Cell 163, 1400–1412 (2015).

15. D. Grun, L. Kester, A. van Oudenaarden, Validation of noise models for singlecell transcriptomics. Nat Methods 11,637–640 (2014).

16. F. Paul et al., Transcriptional Heterogeneity and Lineage Commitment in Myeloid Progenitors. Cell 163, 1663–1677 (2015).

17. S. A. Mani et al., The epithelial-mesenchymal transition generates cells with properties of stem cells. Cell 133, 704–715 (2008).

18. D. A. Jaitin et al., Massively parallel single-cell RNA-seq for marker-free decomposition of tissues into cell types. Science 343, 776–779 (2014).

19. M. B. Eisen, P. T. Spellman, P. O. Brown, D. Botstein, Cluster analysis and display of genome-wide expression patterns. Proceedings of the National Academy of Sciences 95, 14863–14868 (1998).

20. E. Segal et al., Module networks: identifying regulatory modules and their condition-specific regulators from gene expression data. Nat Genet 34, 166–176 (2003).

21. L. H. Hartwell, J. J. Hopfield, S. Leibler, A. W. Murray, From molecular to modular cell biology. Nature 402, C47–52 (1999).

22. A. D. Amir el et al., viSNE enables visualization of high dimensional single-cell data and reveals phenotypic heterogeneity of leukemia. Nat Biotechnol 31, 545-552 (2013).

23. J. H. Levine et al., Data-Driven Phenotypic Dissection of AML Reveals Progenitor-like Cells that Correlate with Prognosis. Cell 162, 184–197 (2015).

24. Q. Li et al., Dynamics inside the cancer cell attractor reveal cell heterogeneity, limits of stability, and escape. Proc Natl Acad Sci U S A 113, 2672–2677 (2016).

25. J. M. Lee. (Springer, 2001).

26. C. Trapnell et al., The dynamics and regulators of cell fate decisions are revealed by pseudotemporal ordering of single cells. Nat Biotechnol 32, 381–386 (2014).

27. R. R. Coifman, S. Lafon, Diffusion maps. Appl Comput Harmon A 21, 5–30 (2006).

28. R. R. Coifman et al., Geometric diffusions as a tool for harmonic analysis and structure definition of data: Diffusion maps. P Natl Acad Sci USA 102, 7426–7431 (2005).

29. R. L. Sparrow, G. Healey, K. A. Patton, M. F. Veale, Red blood cell age determines the impact of storage and leukocyte burden on cell adhesion molecules, glycophorin A and the release of annexin V. Transfusion and apheresis science 34, 15–23 (2006).

30. S. Lamouille, J. Xu, R. Derynck, Molecular mechanisms of epithelial-mesenchymal transition. Nat Rev Mol Cell Biol 15, 178–196 (2014).

31. N. Tiwari et al., Sox4 is a master regulator of epithelial-mesenchymal transition by controlling Ezh2 expression and epigenetic reprogramming. Cancer Cell 23, 768–783 (2013).

32. W. M. Rand, Objective criteria for the evaluation of clustering methods. Journal of the American Statistical association 66, 846–850 (1971).

33. R. R. Coifman, S. Lafon, Geometric harmonics: A novel tool for multiscale out-of-sample extension of empirical functions. Appl Comput Harmon A 21, 31–52 (2005).

34. P. Grassberger, I. Procaccia, Characterization of strange attractors. Physical review letters 50, 346 (1983).

35. M. Francesconi, B. Lehner, The effects of genetic variation on gene expression dynamics during development. Nature 505, 208–211 (2014).

36. M. N. Arbeitman et al., Gene expression during the life cycle of Drosophila melanogaster. Science 297, 2270–2275 (2002).

37. Z. I. Botev, J. F. Grotowski, D. P. Kroese, Kernel Density Estimation Via Diffusion. Ann Stat 38, 2916–2957 (2010).

38. K. Sricharan, R. Raich, A. O. Hero, Estimation of nonlinear functionals of densities with confidence. IEEE Transactions on Information Theory 58, 4135-4159 (2012).

39. M. Rosenblatt, Remarks on Some Nonparametric Estimates of a Density-Function. Ann Math Stat 27, 832–837 (1956).

40. E. Parzen, Estimation of a Probability Density-Function and Mode. Ann Math Stat 33, 1065–& (1962).

41. D. W. Scott, Multivariate density estimation: theory, practice, and visualization. (John Wiley & Sons, 2015).

42. K. Aigner et al., The transcription factor ZEB1 δEF1) promotes tumour cell dedifferentiation by repressing master regulators of epithelial polarity. Oncogene 26, 6979–6988 (2007).

43. D. Achlioptas, F. McSherry, Fast computation of low-rank matrix approximations. Journal of the ACM (JACM) 54, 9 (2007).

44. E. Candes, B. Recht, Exact matrix completion via convex optimization. Communications of the ACM 55, 111–119 (2012).

45. Y. a. Kang et al., SMAD4 regulates cell motility through transcription of N-cadherin in human pancreatic ductal epithelium. PloS one 9, e107948 (2014).

46. B. Adamson et al., A multiplexed single-cell CRISPR screening platform enables systematic dissection of the unfolded protein response. Cell 167, 1867–1882. e1821 (2016).

47. P. G. de Lena, A. Paz-Gallardo, J. M. Paramio, R. Garcia-Escudero, Clusterization in head and neck squamous carcinomas based on lncRNA expression: molecular and clinical correlates. bioRxiv, 105999 (2017).

48. A. Dixit et al., Perturb-seq: dissecting molecular circuits with scalable single-cell RNA profiling of pooled genetic screens. Cell 167, 1853–1866 e1817. (2016).

49. R. Zilionis et al., Single-cell barcoding and sequencing using droplet microfluidics. Nature Protocols 12, 44–73 (2017).

